# Identification of a Pathway for Electron Uptake in *Shewanella oneidensis*

**DOI:** 10.1101/2021.01.12.426419

**Authors:** Annette R. Rowe, Farshid Salimijazi, Leah Trutschel, Joshua Sackett, Oluwakemi Adesina, Isao Anzai, Liat H. Kugelmass, Michael H. Baym, Buz Barstow

## Abstract

Extracellular electron transfer (EET) could enable electron uptake into microbial metabolism for the synthesis of complex, energy dense organic molecules from CO_2_ and renewable electricity^1-6^ . EET could do this with an efficiency comparable to H_2_ -oxidation^7,8^ but without the need for a volatile intermediate and the problems it causes for scale up^9^. However, naturally occurring electroactive organisms suffer from a number of technical drawbacks. Correcting these will require extensive knowledge of the genetics and mechanisms of electron uptake. To date, studies of electron uptake in electroactive microbes have focused on shared molecular machinery also used for anaerobic mineral reduction, like the Mtr EET complex in the electroactive microbe *Shewanella oneidensis* MR-1^10-14^. However, this shared machinery cannot explain all features of electron uptake, hindering efforts to engineer electron uptake. To address this, we screened gene disruption mutants for 3,667 genes, representing ≈ 99% of all non-essential genes, from the *S. oneidensis* whole genome knockout collection using a redox dye oxidation assay as a proxy for electron uptake. Confirmation of electron uptake using electrochemical testing allowed us to identify five genes from *S. oneidensis* that are indispensable for electron uptake from a cathode. Knockout of each gene eliminates extracellular electron uptake, yet in 4 of the 5 cases produces no significant defect in electron donation to an anode, highlighting a distinct role for these loci in electron uptake vs. donation. This result highlights an electronic connection between aerobic and anaerobic electron transport chains that allow electrons from the reversible EET machinery to be coupled to different respiratory processes in *S. oneidensis*. Furthermore, we find homologs to these genes across many different genera suggesting that electron uptake by EET coupled to respiration could be a widespread phenomenon. These gene discoveries provide a foundation for studying this phenotype in exotic metal-oxidizing autotrophic microbes. Additionally, we anticipate that the characterization of these genes will allow for the genetic improvement of electron uptake in *S. oneidensis*; and genetically engineering electron uptake into a highly tractable host like *E. coli* to complement recent advances in synthetic CO_2_ fixation^15^.

## Introduction

Electromicrobial production technologies aim to combine the flexibility of CO_2_ -fixing and C_1_ -assimilating microbial metabolism for the synthesis of complex, energy dense organic molecules from CO_2_ and renewable electricity^1-6^ . Already, the Bionic Leaf device has demonstrated that technologies of this class could dramatically exceed the efficiency of photosynthesis^7,8^. However, while highly efficient at lab scale, the Bionic Leaf relies on H_2_ -oxidation to transfer electrons from the electrode to microbes, and the low solubility of H_2_ in water would pose a significant challenge for scale-up of this and related technologies^9^.

Extracellular electron uptake (EEU) as an electron source for metabolism could allow engineers to circumvent the scale-up limitations of H_2_ -oxidation. Naturally occurring electroactive microbes can produce acetate and butyrate from CO_2_ and electricity with Faradaic efficiencies exceeding 90%^16^. Furthermore, theoretical analysis suggests that the upper limit efficiency of electromicrobial production of biofuels by EEU could rival that of H_2_ -mediated systems^9^. However, naturally occurring electroactive organisms capable of EEU suffer from multiple technical drawbacks. Most notably they have a low-tolerance to high-osmotic strength electrolytes, requiring the use of electrolytes that confer low electrochemical cell conductivity and thus a low overall energy efficiency. Additionally, they have a poor ability to direct metabolic flux to a single product more complex than acetate or butyrate^16^. Correcting these problems to take full advantage of EEU”s potential by genetic engineering^17^ will require extensive knowledge of the genetics of EEU.

Growing evidence suggests that the model electroactive microbe *S. oneidensis* can couple extracellular electron uptake (EEU) to the regeneration of ATP and NADH, both essential precursors for biosynthesis^13^, by reversal of its Extracellular Electron Transfer (EET) pathway (**Fig. 1**), making it an attractive chassis organism for electromicrobial production. However, EEU machinery in *S. oneidensis* appears to involve more than just operating the well-characterized EET machinery in reverse^13,18^. EEU in *S. oneidensis* can link cathodic current with multiple terminal electron acceptors, including oxygen, which draws into question how electrons transfer between canonically discrete electron transport chains. Finding this machinery has been hindered by the lack of high-throughput assays for electron uptake and the challenge of developing screens for non-growth-related phenotypes. Even with recent advances in high-throughput electrode arrays^19^, searching through the thousands of genes in even a single microbial genome by direct electrochemical measurements remains impractical.

**Figure 1.**
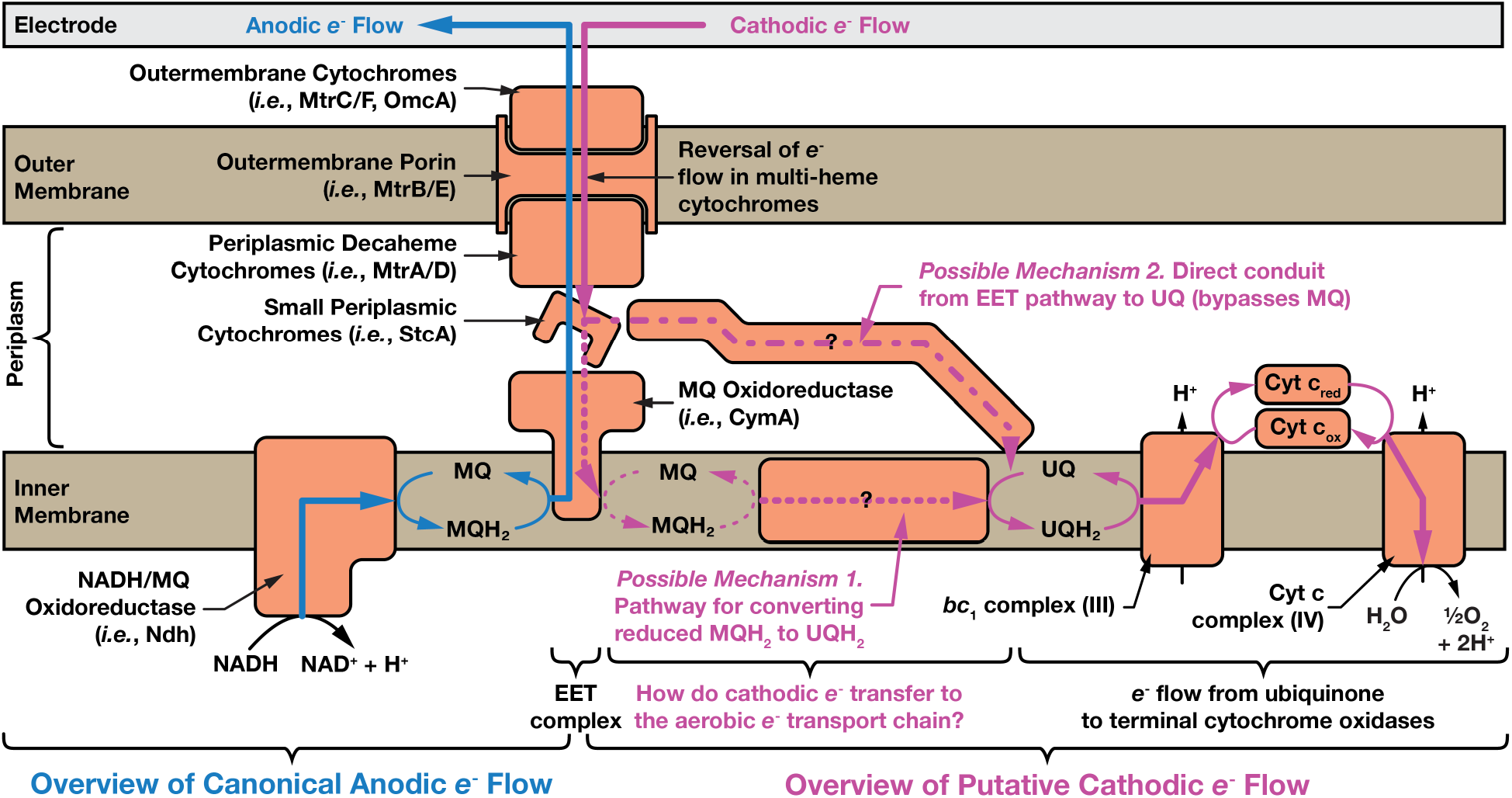
Electron uptake in the model electroactive microbe *Shewanella oneidensis* MR-1 cannot be fully explained by reversal of its extracellular electron transfer pathway. The canonical anodic extracellular electron transport (EET) pathway for electron deposition is shown in light blue, and the putative cathodic extracellular electron uptake (EEU) pathway is shown in pink. Known electron transfer pathways are denoted with solid lines, while speculated transfer pathways are shown as dashed lines. Two possible mechanisms for transfer of cathodic electrons from the Mtr EET complex to the ubiquinone pool and onto terminal cytochrome oxidases are highlighted.

To address this, we developed a rapid colorimetric assay to screen all 3,667 members of the *S. oneidensis* whole genome knockout collection^20,21^ and characterize the genetics of EEU. The assay relies upon oxidation of the reduced form of the redox dye anthra(hydro)quinone-2,6-disulfonate (AHDS_red_ for the reduced form and AQDS_ox_ for the oxidized form) and is coupled to reduction of the anaerobic terminal electron acceptors fumarate and nitrate^22-24^ (**Fig. 2**). While AHDS_red_ /AQDS_ox_ redox dye assays are not a perfect proxy for EEU and EET, they are capable of identifying many components of the *S. oneidensis* EET machinery^20^. Simplified AHDS_red_ oxidation time courses for assay controls and wild-type *S. oneidensis* are shown in **Fig. 2B** along with a detailed time course in **Fig. S1**.

**Figure 2.**
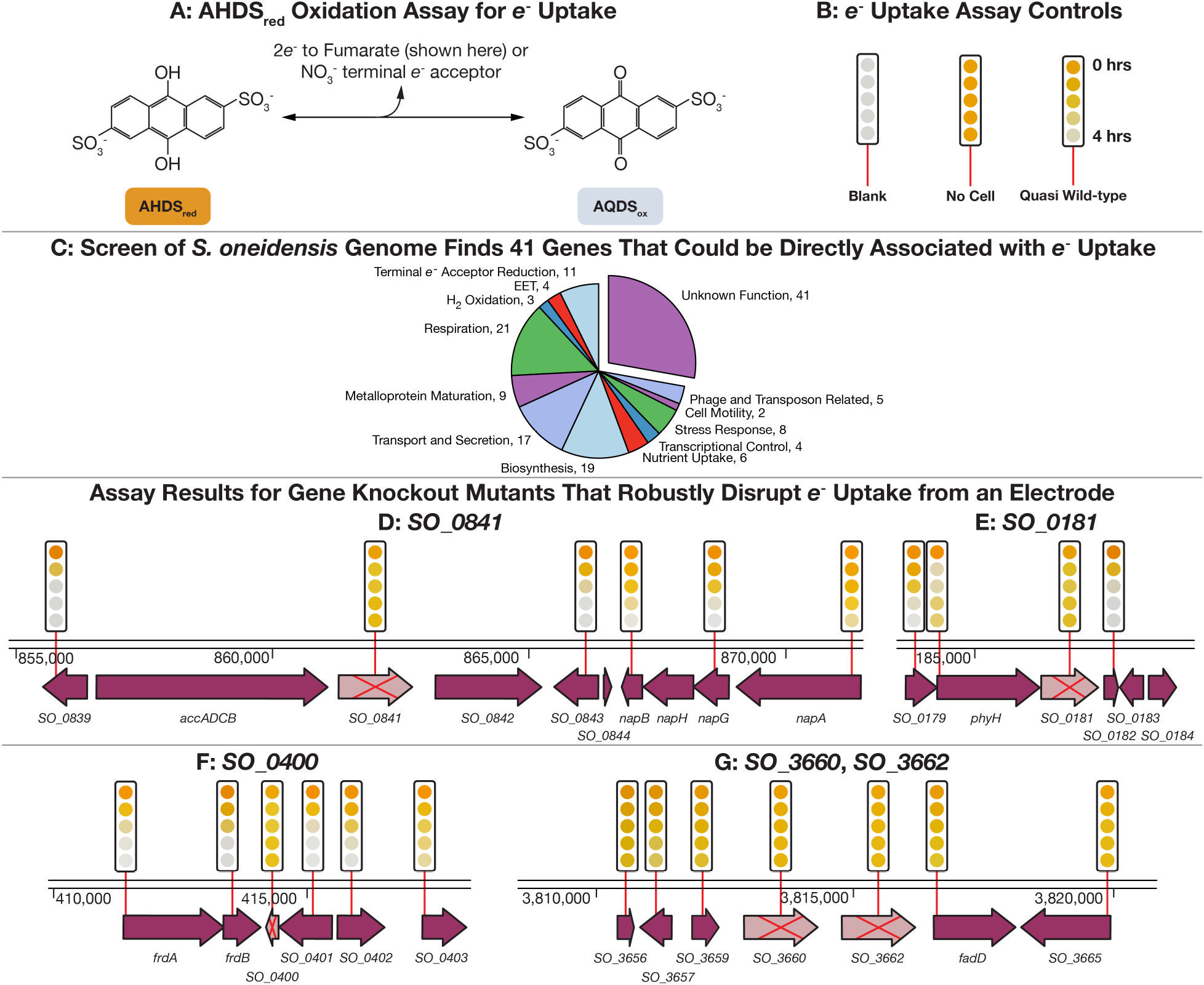
A genome-wide screen of *S. oneidensis* finds 150 genes that disrupt electron uptake. All 3,712 members of the *S. oneidensis* whole genome knockout collection were screened for electron uptake capability with an AHDS_red_ oxidation assay. (**A**) AHDS_red_/AQDS_ox_ redox reaction is used as a proxy for extracellular electron uptake. AHDS_red_ changes color from orange to clear when oxidized. Electrons are transferred to either a fumarate or nitrate terminal electron acceptor by *S. oneidensis*. Data shown uses fumarate. (**B**) Blank, no-cell and quasi-wild-type (transposon mutants that contain a kanamycin cassette but have no effect on AHDS_red_ oxidation) controls. The color of the AHDS_red_ dye is recorded photographically and displayed at 1 hour intervals after the start of the experiment by a series of colored circles above each gene. Further information on this assay can be seen in **Fig. S1** and **Materials and Methods**. (**C**) The electron uptake assay associates 150 genes with electron uptake. Electron uptake failure can be explained in 109 cases, but in 41 cases it fails for unknown reasons, implicating these genes in a novel electron uptake process. Full screening results and functional categorizations are shown in **Table S1**. (**D** to **G**) AHDS_red_ oxidation assay results for deletion mutants containing knockouts of genes highlighted in this article (pink arrow with a red cross through the center) along with gene disruption mutants for surrounding genes (purple arrow, with a red line indicating the location of the transposon insertion).

To ensure that genes identified by high-throughput screening are involved in EEU with solid surfaces, a subset were tested in electrochemical systems (**Fig. 3**): the gold standard for measuring EEU^25,26^.

**Figure 3.**
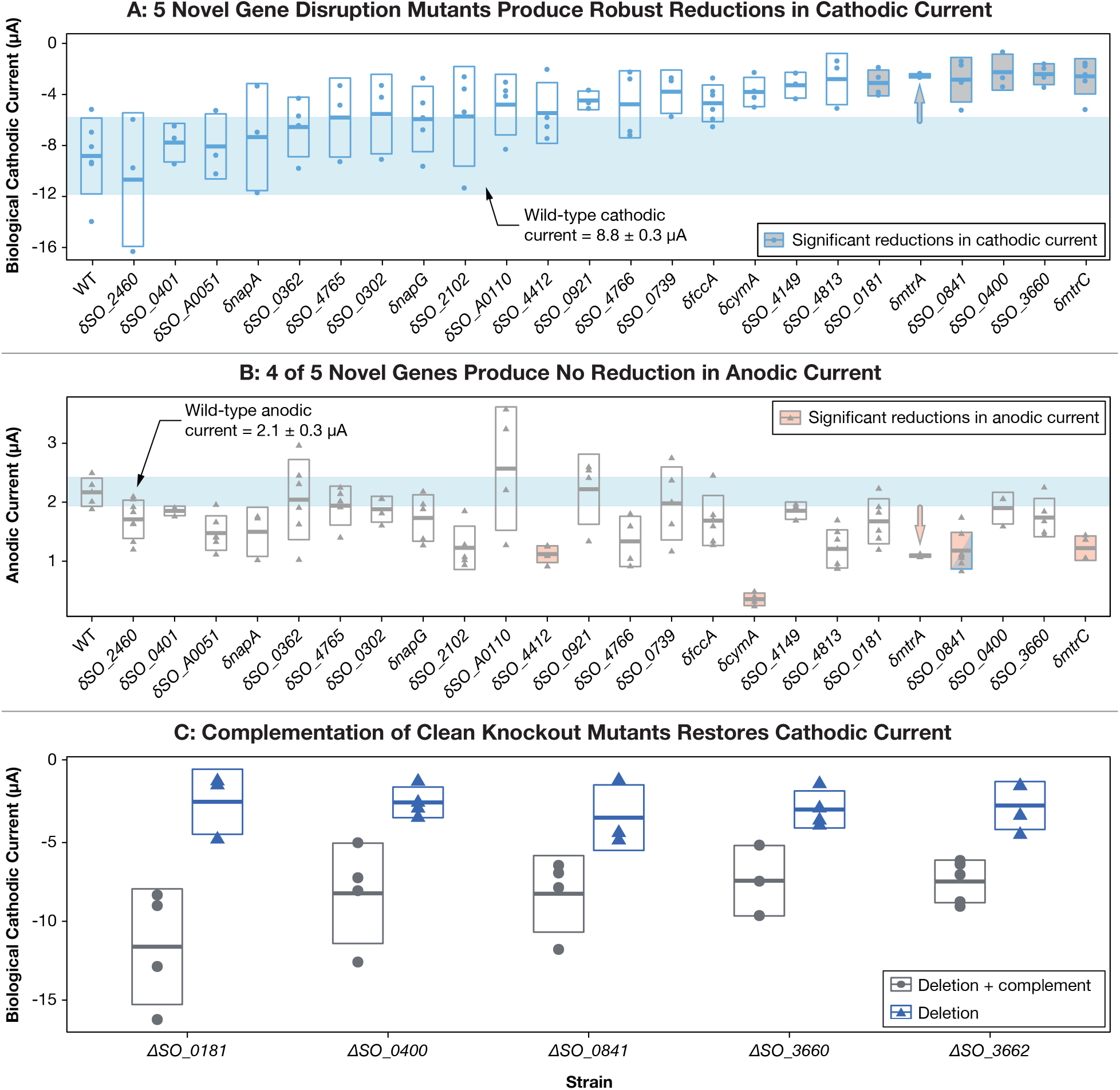
Electrochemical measurements confirm robust extracellular electron uptake phenotype for 5 novel *S. oneidensis* mutants identified by our high-throughput screen. (**A**) We measured the cathodic biological current for 18 *S. oneidensis* mutants that produced AHDS_red_ oxidation failure for unknown reasons, control mutants and wild-type (WT). ANOVA results indicated that there was a significant variation in means of the strains from one another (F-value = 5.94; Pr(>F) = 2.34 × 10^−9^), while individual comparisons specifically revealed *δSO_0181, δmtrA, δSO_0841, δSO_0400, δSO_3660* and *δmtrC* (*δ* indicates a transposon insertion mutant) to all display significantly lower cathodic currents (*p* < 0.05) than WT . (**B**) However, 4 of the 5 robust electron uptake mutants do not have a significant effect on anodic current production (electron deposition), the well-characterized phenotype of *S. oneidensis*. Analysis of anodic current values (F-value= 5.801, Pr(>F)=2.10 × 10^−10^) revealed that of the 5 novel mutants, only *δSO_0841* shows anodic current production different from WT (p < 0.05). (**C**) Gene deletion (indicated by *Δ)* confirms the electron uptake phenotype of disruption mutants, and complementation of the deleted gene restores electron uptake phenotype. Further information on electrochemical methods can be seen in **Materials and Methods**

## Results and Discussion

### High-throughput Electron Uptake Screen Finds 41 Genes with Unknown Function

We identified mutants in 150 coding and intergenic regions in the *S. oneidensis* genome that slowed or eliminated AHDS_red_ oxidation (**Table S1**). 109 of these mutants were grouped by gene annotation into functional categories that satisfactorily explain the slowing or failure of AHDS_red_ oxidation (**Fig. 2C**). For example, disruption of the periplasmic fumarate reductase (*δfccA*; we refer to transposon disruption mutants with *δ*, and gene deletion mutants with *Δ*) eliminates AHDS_red_ oxidation when using fumarate as a terminal electron acceptor. Detailed time courses of AHDS_red_ oxidation for selected anticipated hits from the genome-wide screen are shown in **Fig. S2**. Of note, 41 of the AHDS_red_ oxidation-deficient mutants could not be assigned to an established functional category, suggesting that their function might be involved in electron uptake (**Fig. 2C**). AHDS_red_ oxidation time courses for knockout mutants where we later observed a cathode phenotype are shown in **Figs. 2D** to **G**, along with those for mutants with disruptions in adjacent genes. Detailed time courses for these mutants are shown in **Fig. S3**.

### Electrochemical Measurements Confirm Robust Extracellular Electron Uptake Phenotype of Five Novel Mutants

We selected 17 of the 41 “unknown function” *S. oneidensis* AHDS_red_ oxidation-deficient mutants for further on-electrode testing. These mutants were chosen for annotations that indicated possible redox activity (*e*.*g*., *δSO_3662*), interaction with the quinone pool (*e*.*g*., *δSO_0362, δSO_0400*), along with mutants with no annotation at all. We also selected 2 positive control mutants (*δmtrA* and *δmtrC*) and 5 negative control mutants (*δcymA, δfccA, δnapA, δnapG*, and *δSO_0401*). While disruption of genes coding for anaerobic terminal reductases (*δfccA, δnapA, δnapG*) register as hits in the AHDS_red_ oxidation assay, these mutants should not disrupt electron uptake from a cathode when O_2_ is used as a terminal electron acceptor. *δSO_0401* was chosen as it is adjacent to a hit (*δSO_0400*) in the AHDS_red_ oxidation assay, but does not itself produce a hit.

Biofilms of each of the mutants were grown on ITO working electrodes in a three-electrode bio-electrochemical system^13^. For analysis of electron uptake, the working electrodes were poised at −278 mV vs. the Standard Hydrogen Electrode (SHE). Significant negative currents (*i*.*e*., electrons flowing from the working electrode to the biofilm/solution) were only observed in the presence of O_2_ as a terminal electron acceptor. To quantify the amount of negative current due to biological vs. non-biological processes, the electron transport chain was inhibited at the end of each experiment with the ubiquinone mimic, Antimycin A and remaining abiotic current was measured (**Fig. S4**). Each mutant was tested in at least three replicate experiments.

Most of the 17 mutants of unknown function demonstrate a limited to modest change in average electron uptake from the working electrode (**Figs. 3A, S5A, S5C, S5D**, and **Table S2**). As expected, mutants that disrupt components of the well-known Mtr EET complex produce significant reductions (*p* value < 0.05) in electron uptake except for *cymA*^10,13^. Though *cymA* was previously shown to be important under anaerobic cathodic conditions^10^, only a small reduction in electron uptake was noted under aerobic conditions, consistent with previous results^13^. It is plausible that the other unknown genes tested that did not generate a cathodic phenotype play a previously uncharacterized role in one of the other subcategories highlighted in the AHDS_red_ assay rather than electron uptake, such as the reduction of fumarate or nitrate, as opposed to O_2_.

Disruption mutants in four genes that code for proteins that are of unknown function (*δSO_0181, δSO_0400, δSO_0841, δSO_3660*) demonstrated a highly significant reduction in electron uptake (*p* value < 0.05) (**Fig. 3A**). To further verify that this reduction was due to the loss of the disrupted gene and not due to any polar effects, we made a clean deletion for each, which all demonstrated a decreased electron uptake phenotype compared to wild-type (**Fig. 3C**). Though *δSO_3662* was not initially tested on cathodes, Δ*SO_3662* (annotated as an inner membrane ferredoxin), was constructed and demonstrated both an AHDS_red_ oxidation phenotype (**Fig. 2G and Fig. S3**) and a cathodic phenotype (**Fig. 3C**). Furthermore, complementation of the knockout mutants with a plasmid encoding the deleted gene restored electron uptake function in all mutants (**Fig. 3C**).

### Four of the Five Novel Genes are Only Involved in Electron Uptake

Electron uptake disruption in these five gene deletion mutants cannot be explained by changes to biofilm production or cell growth rate (**Table S3**). No statistically significant difference in on-electrode protein abundance was seen for any of the mutants tested (**Table S2**). Furthermore, no statistically significant difference in growth rate between wild-type *S. oneidensis* and any mutant was observed in minimal media under aerobic or anaerobic conditions with the exception of a longer lag time observed for complementation mutants containing an extra plasmid and grown in the presence of chloramphenicol and/ or kanamycin (**Table S3** and **Fig. S6**).

In addition to electron uptake, we also analyzed the extracellular electron deposition of each of the 24 mutants selected for on-electrode testing (**Fig. 3B**). Of the 5 novel EEU genes identified here (*SO_0181, SO_0400, SO_0841, SO_3660*, and *SO_3662)*, only disruption of *SO_0841* significantly reduces both electron uptake and deposition. The *δSO_0841* disruption mutant produces half the positive current of wild-type *S. oneidensis*, similar to the effect of disruption of *mtrA* and *mtrC* (**Table S2** and **Fig. 3B**). Interestingly, no growth phenotype was observed for cells grown on soluble (iron citrate) or insoluble iron (iron oxyhydroxide), suggesting that SO_0841 does not play a significant role in EET to minerals (data not shown).

### Sequence Analysis Suggests Gene Functions and Shows Electron Uptake Genes are Widely Distributed Across Species

Phylogenetic analysis of the five novel EEU genes identified here suggests they are broadly conserved across *Shewanella* species, and across numerous clades of the *Gammaproteobacteria*.

### SO_0841

Sequence analysis suggests that SO_0841 is involved in cell signaling. *SO_0841* encodes a transmembrane protein with 250 amino acid long periplasmic region and a 250 amino acid long cytoplasmic domain containing a GGDEF c-di-GMP signaling domain. As a phenotype was only observed on electrodes, and not iron minerals, the role of this protein in traditional EET remains unclear. The GGDEF domain is often used for regulation of biofilm formation and cell motility^27^, however disruption of *SO_0841* has on impact on biofilm formation. This suggests a more specific role for SO_0841 in on-electrode EET. *SO_0841* also has a broad distribution of homologs ranging across the *Proteobacteria* (**Fig. S7)**. Homologs of *SO_0841* are found in electrochemically active microbes with the capacity for EEU, including *Mariprofundus ferrooxydans*^28^, *Idiomarina loihiensis*^29^ and *Marinobacter* species^29^, which supports a conserved role for this gene in electron uptake.

The remaining four novel genes identified all play significant roles in electron uptake but have no detectable role in electron deposition (**Figs. 3A** and **3B**).

### SO_0181

SO_0181 is predicted to be membrane associated and contains a putative nucleoside triphosphate binding and/or hydrolase domain (suggesting that it interacts with ATP or GTP). Furthermore, *SO_0181* is located immediately upstream of *phyH* which encodes an uncharacterized putative oxidoreductase (part of a family of bacterial dioxygenases with unconfirmed activity in *S. oneidensis*), which also demonstrated an electron uptake phenotype in the AHDS_red_ oxidation screen (**Table S1** and **Fig. 2E**).

There is little bioinformatic support for a direct role of SO_0181 in redox chemistry. However, a role associated with activating or modifying the other redox active proteins involved in EEU seems feasible but needs to be further investigated.

A phylogenetic tree constructed from the 200 closest relatives of *SO_0181* shows a distinct clade specific to the *Shewanellae* (90-100% sequence identity) (**Fig. S8)**. Closely related clades of *SO_0181* are found in the *Pseudomonas, Cellvibrio*, and *Hahella* genera of the *Gammaproteobacteria*, though homologous gene clusters are also seen in *Beta-(Delftia*, and *Acidovorax)* and *Delta-proteobacterial (Archangium*, and *Cystobacter)* lineages. Notably, several of the genera with close homologs of *SO_0181*, including *Delftia* and *Pseudomonas*, have EET-capable representatives within them.

### SO_0400

SO_0400 belongs to a super-family of quinol-interacting dimeric monooxygenses (dimeric α-β barrel superfamily SSF54909). Of proteins of known function, SO_0400 is most closely related to the YgiN quinol monooxygenase in *E. coli*^30^. Structural analysis of YgiN suggests that it interacts with the semiquinone state of quinols and suggests the existence of a novel quinone redox cycle in *E. coli*^30^. Additionally, deletion of *ygiN* in *E. coli* inhibits the transition between aerobic and anaerobic growth^31^. Interestingly, deletion of *SO_0400* does not inhibit the transition between aerobic and anaerobic growth conditions (and vice versa) in *S. oneidensis* (**Fig. S6**). Furthermore, deletion of *SO_0400* did not affect the sensitivity of *S. oneidensis* to oxygen free radicals using a disc diffusion assay with hydrogen peroxide (data not shown). These data suggest a possible novel function in this quinol monooxygenase.

*SO_0400* has a very broad distribution of close homologs found in the *Proteobacteria*, Gram positive *Actinobacteria, Bacteriodetes*, and Archaeal *Methanobrevibacter* (**Fig. S9)**. Within the *Shewanellae* homologs of this quinol monooxygenase are both broadly distributed and tightly clustered in a highly conserved clade, with many homologs exhibiting 95-100% amino acid sequence identity. This may speak to the highly conserved function of this gene within the *Shewanellae* that is possibly distinct from the other homologs observed in other genera.

### SO_3660, SO_3662 and an Electron Uptake Operon

The AHDS_red_ oxidation screen points to the existence of an electron uptake operon in *S. oneidensis*. Disruption of any of the loci from *SO_3656* to *SO_3665* cause failure of AHDS_red_ oxidation (**Fig. 2G**). This putative operon is possibly regulated by *SO_3660*, annotated as a transcriptional regulator. *SO_3662* is annotated as an inner membrane ferredoxin, strongly suggesting a direct role in electron transfer. In-frame deletions of the genes coding for SO_3660 and the putative inner membrane bound ferredoxin *SO_3662* both disrupt electron uptake (**Fig. 3C**), but not deposition.

Phylogenetic trees constructed from the 200 closest homologs of *SO_3600* and *SO_3662* revealed these genes are highly conserved in *Gammaproteobacteria* and across numerous (≈100) *Shewanella* species (**Figs. S10** and **S11**). *SO_3662* appears to be highly conserved among the Order *Alteromodales* in particular.

### Role of Electron Uptake Genes in *S. oneidensis*

Like many other facultative microorganisms including *E. coli*^32^, *S. oneidensis* employs discrete anaerobic and aerobic electron transport chains. Menaquinone is the dominant quinone used by *S. oneidensis* under anaerobic conditions where EET is used for mineral respiration^23^. Conversely, under aerobic conditions ubiquinone is the dominant quinone used by *S. oneidensis*^33.^ Furthermore, ubiquinone is important for reverse electron flow to NADH mediated by Complex I (a NADH:ubiquinone oxidoreductase) under cathodic conditions^13^.

When taken together, the normal isolation of aerobic and anaerobic electron transport chains and the ability of *S. oneidensis* to couple reversal of the anaerobic EET pathway to O_2_ reduction suggests that a specific connection between the two transport chains is likely to exist. However, to our knowledge, the mechanism allowing organisms to transition between one electron transport chain, or quinone pool to another are poorly understood.

We outline two possible mechanisms for a connection between the anaerobic EET pathway and the aerobic electron transport chain in EEU in **Fig. 1**. First, the EET complex could transfer electrons to CymA, which then reduces menaquinone. Electrons could then be transferred from menaquinone to ubiquinone and into the aerobic electron transport chain, finally arriving at Complex IV where they reduce O_2_. This option seems intuitive as under anaerobic conditions using fumarate as an electron acceptor, *S. oneidensis* was shown to require menaquinone in addition to several components of the EET complex to uptake cathodic currents^10^. However, knockout of the gene coding for *cymA* does not disrupt cathodic electron uptake when using O_2_ as a terminal electron acceptor^13^.

The lack of involvement of CymA in cathodic electron uptake under aerobic conditions suggests a second option (**Fig. 1**): that cathodic electrons bypass the menaquinone pool under aerobic conditions.

We speculate that the putative quinol-interacting protein SO_0400 and the putative ferredoxin SO_3662 (and possibly proteins coded by nearby genes) are directly involved in connecting the reverse EET pathway during electron uptake and the aerobic electron transport chain. Notably, the lack of a phenotype for most of these proteins under anodic conditions supports the hypothesis of a distinct connection between a subset of EET proteins and the aerobic electron transport chain during EEU (**Fig. 1**). This work has also highlighted some genes potentially involved in cell signaling or transcriptional responses that may help aid in facilitating electron uptake under specific conditions (*SO_0841, SO_0181*, and *SO_3660*).

Though the motivation of this work stemmed from the application of *S. oneidensis* to electrosynthesis, it is likely that the process of extracellular electron uptake plays a role in *Shewanella* physiology and ecology. It has been shown that minerals in nature can serve a capacitive or electron storing function for microbes^34^. While electron deposition and electron uptake from minerals such as magnetite were shown to function as both sinks and sources for different metabolisms, it is feasible that a single organism with both functionalities could utilize minerals in this way--functionally storing charge akin to a battery.

Though iron-oxidation has only been demonstrated in a single *Shewanella* strain^35^, this could be due to the challenge of distinguishing biologic and abiotic iron oxidation in the absence of growth. As *Shewanella* are not generally capable of carbon fixation, the process of extracellular electron uptake is unlikely to have evolved as a growth linked metabolism. However, previous work has linked electron uptake to maintenance of cell biomass or decreasing the rate of cell death, which could suggest a role in allowing cells to conserve energy under non-growth conditions^13^. Interestingly, these genes appear to be widely conserved across the *Shewanella*, as well as other marine *Gammaproteobacteria* (several of which have also been implicated in extracellular electron uptake). This supports the yet unexplored ecological role for extracellular electron uptake in sediment and/or marine microbes, though our knowledge of the specific activity and role of this process is still at its inception.

## Conclusions

EEU holds significant potential for conversion of CO_2_ and renewable electricity to complex organic molecules. While potential of novel phenotypes like this can be limited by a lack of genetic understanding, especially in non-model organisms, synthetic biology can greatly expand the possibility for their improvement and application.

We used a whole-genome knockout collection previously built with the rapid, low-cost Knockout Sudoku method to screen the *S. oneidensis* genome for redox dye oxidation, a proxy for electron uptake. In this work we have performed detailed electrochemical analyses, focusing on genes encoding proteins of unknown function. We have identified five novel genes in *S. oneidensis* that are involved in extracellular electron uptake from both solid phase and extracellular donors, coupled to both aerobic and anaerobic terminal electron acceptors.

Identification of these important genes lays the foundation for further genetic characterization of metal oxidation in nature, improvement of EEU in *S. oneidensis*, and for synthetically engineering an electron uptake pathway into easily engineered or synthetic microbes for powering recent advances in synthetic CO_2_ fixation^15^ and EET^36^ in *E. coli*.

## Materials and Methods

### Genome-wide AHDS_red_ Oxidation Screen

The *S. oneidensis* whole genome knockout collection^20,21^ was screened for members unable to oxidize the redox dye anthra(hydro)quinone-2,6-disulfonate (AHDS_red_ for the reduced form and AQDS_ox_ for the oxidized form)^23,24^, and subsequently reduce either fumarate^22^ or nitrate.

### Knockout Collection Construction

The *S. oneidensis* whole genome knockout collection was previously built with the Knockout Sudoku whole genome knockout collection construction procedure^20,21^. Prior to high-throughput screening the mutant collection was duplicated with a multi-blot replicator (Catalog Number VP 407, V&P Scientific, San Diego CA, USA) into 96-well polypropylene plates containing 100 µL of LB with 30 µg mL^-1^ kanamycin per well. The plates were labeled with barcodes and registration marks for identification in high-throughput analysis. The plates were sealed with an air porous membrane (Aeraseal, Catalog Number BS-25, Excel Scientific) and grown to saturation overnight (at least 24 hours) at 30°C with shaking at 900 rpm.

### AHDS_red_ Preparation

Solutions of 25 mM AHDS_red_for screening experiments were prepa_red_ electrochemically. 200 to 1,000 mL batches of 25 mM AQDS_ox_were prepared by dissolving 10.307 mg AQDS_ox_powder (Catalog no. A0308, TCI America) per 1 mL of deionized water at 60 °C. The AQDS_ox_ solution was then transferred to a three-electrode electrochemical system (Catalog no. MF-1056, BASI Bulk Electrolysis Cell) inside a vinyl anaerobic chamber (97% N_2_, 3% H_2_, < 20 ppm O_2_; Coy Laboratory Products, Grass Lake MI, USA). The system uses a Ag/AgCl reference electrode, a coiled Pt wire counter electrode inside a fritted counter electrode chamber, and a reticulated vitreous carbon working electrode. The working electrode potential was maintained at 700 mV vs. Ag/AgCl with a digitally controlled potentiostat (PalmSens, EmStat3). To enhance cell conductivity, 3M H_2_SO_4_ acid was added to the working electrode, allowing the cell current to rise to ≈ 2 mA. AHDS_red_ reduction is assumed to be complete when the solution is translucent red in color, typically after ≈ 7 hours. The pH of the AHDS_red_ stock solution was returned to 7.2 by addition of 3M NaOH.

### Assay Media Preparation

*Shewanella* Basal Media (SBM) was used for all AHDSred oxidation assays. SBM consists of: ammonium chloride (NH_4_Cl) (0.0086 M); dibasic potassium phosphate (K_2_HPO_4_) (0.0013 M); monobasic potassium phosphate, (KH_2_PO_4_) (0.0017 M); magnesium sulfate heptahydrate (MgSO_4_.7H_2_O) (0.0005 M); ammonium sulfate (NH4)_2_SO_4_) (0.0017 M); and HEPES (0.1 M). The media was buffered to pH 7.2 with sodium hydroxide (NaOH) and sterilized by autoclave. Trace mineral supplement (5 mL L^-1^; Catalog no. MD-TMS, American Type Culture Collection (ATCC), Manassas, VA, USA) and vitamin supplement (5 mL L^-1^; Catalog no. MD-VS, ATCC) were added aseptically.

Solutions of 25 mM potassium nitrate and 25 mM sodium fumarate adjusted to pH 7.3 and filter sterilized were prepared as terminal electron acceptors for the AHDS_red_ oxidation screens. These solutions were transferred to an anaerobic chamber at least the evening prior to any experiment to allow de-oxygenation.

### AHDS_red_ Oxidation Screens with Nitrate

Aliquots of 10 µL of culture of each mutant from the replicated *Shewanella* whole genome knockout collection were transferred to 96-well polystyrene assay plates filled with 50 µL of SBM using a 96-channel pipettor (Liquidator 96, Mettler-Toledo Rainin LLC, Oakland, CA, USA). The assay plates were transferred to an anaerobic chamber for de-oxygenation for at least 9 hours.

Following de-oxygenation, 40 µL of 1:1 mixture of 25 mM AHDS_red_ and 25 mM potassium nitrate were added to each well of the assay plates with a multi-channel pipettor. The final concentration of both AHDS_red_ and potassium nitrate in each well was 5 mM. Each assay plate was immediately photographed after media addition with the macroscope. All plates were repeatedly imaged for at least ≈ 40 hours. After completion of the experiment, photographs were analyzed with the macroscope image analysis software.

### AHDS_red_Oxidation Screen with Fumarate

Aliquots of 1 µL of culture of each mutant from the replicated *S. oneidensis* whole genome knockout collection were transferred to 96-well polystyrene assay plates filled with 59 µL of SBM using a multi-blot replicator (Catalog Number VP 407, V&P Scientific, San Diego CA, USA). The assay plates were transferred to an anaerobic chamber for de-oxygenation for at least 9 hours. AHDS_red_ oxidation was found to proceed much faster with fumarate than with nitrate, so the volume of cells added to the assay was reduced to make data collection more manageable.

Following de-oxygenation, 40 µL of 1:1 mixture of 25 mM AHDS_red_ and 25 mM sodium fumarate were added to each well of the assay plates with a multi-channel pipettor. The final concentration of both AHDS_red_ and sodium fumarate in each well was 5 mM. Each assay plate was immediately photographed after media addition with the macroscope. All plates were repeatedly imaged for at least ≈ 40 hours. After completion of the experiment, photographs were analyzed with the macroscope image analysis software.

| *AHDS<sub>red</sub> Oxidation with Nitrate Assay* |  |  | *AHDS<sub>red</sub> Oxidation with Fumarate Assay* |  |  |
| --- | --- | --- | --- | --- | --- |
| Component | Volume (μL) | Final Conc. (mM) | Component | Volume (μL) | Final Conc. (mM) |
| *Shewanella* Basal Media | 50 | - | *Shewanella* Basal Media | 59 | - |
| Saturated Culture | 10 | - | Saturated Culture | 1 | - |
| 25 mM KNO<sub>3</sub> | 20 | 5 | 25 mM Na Fumarate | 20 | 5 |
| 25 mM AHDS<sub>red</sub> | 20 | 5 | 25 mM AHDS<sub>red</sub> | 20 | 5 |
| **Total Volume (μL)** | 100 |  | **Total Volume (μL)** | 100 |  |

### Macroscope Data Acquisition System

An automated photographic data acquisition system was used to record the results of AHDS_red_ oxidation assays. The device consists of a digital single lens reflex (DSLR) camera (any member of the Canon Rebel Series) controlled by a macOS computer running a custom data acquisition program that downloads images to the computer and timestamps them. The camera shutter is controlled by a single-switch foot pedal (vP-2, vPedal), leaving both hands free to manipulate 96-well plates. The camera is mounted to a frame constructed with extruded aluminum rails (T-slot). Barcodes attached to each plate enable images to be automatically sorted. Registration marks on the barcodes allow for identification of well positions and each well to be associated with a specific gene knockout mutant. The device allows a stack of 200 96-well plates to be imaged in ≈ 15 minutes. This process can be repeated immediately, allowing each plate to be quasi-continuously imaged.

This work has used two variants of the device: the first to image AHDS_red_ oxidation in transparent 96-well assay plates, and the second to measure bacterial growth in 96-well storage plates covered with an air porous membrane (Aeraseal, Catalog Number BS-25, Excel Scientific). In the first configuration, the camera is mounted above the AHDS_red_ oxidation assay plates. Each plate is placed inside a laser-cut acrylic holder and illuminated by an LED light pad from below (A920, Artograph). Barcodes are printed on transparent labels (Catalog no. 5660, Avery) and attached to the top of each plate. In the second configuration, the camera is mounted below the plate and illumination is provided from above with an LED light pad. A white barcode (SIDE-1000, Diversified Biotech, Dedham, MA, USA) is attached to the side of the plate and viewed through a 45° right-angle mirror (Catalog no. 47-307, Edmund Optics, Barrington, NJ, USA).

### Analysis of Macroscope Images

A custom Macroscope Image Analyzer program was developed with Pyzbar^37^, Pillow^38^, Numpy^39^, Matplotlib^40^, and OpenCV-Python^41^ using Python3. The program was used to process images taken with the macroscope device and aided in detection of loss of function mutants. The program handled the image analysis in four steps: creating barcodes, organizing the images collected with the macroscope by barcodes, collecting the data from the images, and finally presenting the data for analysis.

An additional image analysis algorithm was developed with SciKit Image^42^ and SimpleCV^43^ to test images of 96-well storage plates for cross-contamination and growth-failure events by comparison with the collection catalog. The image analysis algorithm updated the record for each well in the collection catalog with growth information to assist in rejection of false positives in the AHDS_red_ oxidation screen due to growth failure.

The Macroscope Image Analyzer software is available on GitHub at https://github.com/barstowlab/macroscope-image-analyzer.

As AHDSred is oxidized to AQDSox, it changes color from yellow-orange to clear. Almost all information on the reduction state of the AHDS_red_/AQDS_ox_ dye can be found in the blue color channel of the assay plate images. At the start of the assay, the intensity of the blue color channel is low, and the dye is orange. As the AHDS_red_/AQDS_ox_ dye is oxidized and becomes clear, the intensity of the red channel remains approximately constant, with a small increase in green channel intensity and a large increase in the blue channel intensity. However, we found that reporting the blue channel intensity as a proxy for the AHDS_red_/AQDS_ox_ redox state to be unintuitive.

To aid the reader and experimenter, we used the RGB color data,

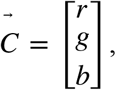

to calculate a single number that represents how ‘yellow’ a well is. The vector overlap (dot product) was calculated between the current color of the well and the most saturated yellow color in the assay photographic data set.

The reference yellow color,

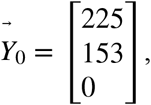

was calculated relative to the reference white color,

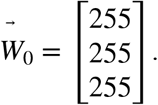

Thus, the transformed yellow reference,

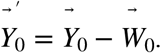

The transformed well color, relative to the white reference,

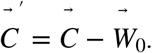

The yellow intensity was calculated by normalizing dot product between the transformed well color and the transformed yellow reference,

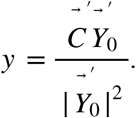

The normalized yellow intensity has a maximum value of 1 when the well is yellow and a minimum value of 0 when it is clear.

The time series of colors for each gene shown as colored circles above each gene in **Fig. 2** were generated by an algorithm that interpolated the multi-replicate average of mean well-center color values for that mutant at 0, 1, 2, 3, and 4 hours after the initiation of the oxidation experiment. A visual explanation of this process is shown in **Fig. S1**. AHDS_red_ oxidation rates reported in **Fig. S5** were calculated by a linear fit to the linear portion of the yellow intensity curve with DATAGRAPH (Visual Data Tools).

### Bio-electrochemical Methods

#### Bio-electrochemical Cell Construction and Experimental Conditions

A 3-electrode electrochemical cell based on a design by Okamoto *et al*.^44^ was assembled in house, with the exception of a salt bridge that was included to contain the reference electrode (Part no. MF-2031, BASi, West Lafeyette, IN, USA). As described, the cell consisted of a working electrode made of ITO-(indium tin oxide) plated glass (Delta Technologies, Ltd., Loveland, CO, USA), a counter electrode of platinum wire, and a Ag/AgCl reference electrode suspended in 1 M KCl (HCH Instruments, Inc., Austin, TX, USA). The reactor volume contained approximately 20 mL of liquid with a working electrode surface area of 10.68 cm^2^.

The electrochemical cell was used for chronoamperometry (CA) experiments that measure change in current over time and cyclic voltammetry (CV) experiments that measure current in response to a change in voltage. Both types of experiment were controlled with a 16-channel potentiostat (Biologic, France). In anodic CA experiments the working electrode was poised at 422 mV vs. SHE (Standard Hydrogen Electrode). In cathodic CA experiments the working electrode was poised at −278 mV. In CV experiments the working electrode potential was scanned between +422 mV and −378 mV vs. the SHE at a rate of 1 mV s^-1^.

#### *Shewanella* Culturing Conditions

Cultures of wild-type *S. oneidensis* and *S. oneidensis* mutants were grown from glycerol stocks overnight in Luria Broth (LB) prior to each experiment. Aerobic and Anaerobic growth curves for each strain were performed in a *Shewanella* Defined Media (SDM)^45^. Aerobic cultures were performed in 50 mL volumes using 10 mM lactate as an electron acceptor shaking at 150 rpm at 30 °C. Kanamycin (Kan) and Chloramphenicol (Chl) were added to LB and SDM media at concentrations of 100 µg mL^-1^ and 34 µg mL^-1^ respectively for selection of transposon (Kan), clean gene deletion (Kan), and complementation (Chl + Kan) strains. The same growth conditions were used for anaerobic growth curves with the exception that the media contained 20 mM fumarate and was purged with nitrogen gas for 10 min in serum vials.

For growth curves, strains were inoculated from overnights at a 100-fold dilution. Optical densities were recorded for triplicate cultures at 600 nm ever 2-3 hours. For cathodic growth, an overnight culture was back-diluted by a factor of 100 into SDM with 10 mM lactate and grown overnight. The overnight cultures grown in SDM were pelleted and resuspended in fresh SDM to an optical density at 600 nm of 0.1 20 mL of the resuspended culture was transferred to the working electrode chamber of an electrochemical reactor. The reactor was attached to 16-channel potentiostat (BioLogic) and the culture was anode conditioned by poising the working electrode at +422 mV vs. SHE^12,13^. Anaerobic conditions needed to encourage biofilm formation and anodic current generation were maintained by continuous purging with N_2_.

After approximately 24 hours the reactors were detached from the potentiostat and the media containing planktonic cells was carefully removed to avoid disturbing the biofilm on the working electrode. The reactor was then refilled with 20 mL of fresh carbon-free SDM^13^. The reactors were then reattached to the potentiostat and the working electrode was cathodically-poised at −378 mV vs. SHE. Air was slowly bubbled into the reactors via an aquarium pump until a steady stream was reached to provide a source of O_2_.

To determine the portion of the cathodic current due to biological processes the respiratory inhibitor Antimycin A was added to the electrochemical cell working electrode chamber to a final concentration of 50 µM. To control for the effects of DMSO (the solvent for Antimycin A), blank injections of DMSO were made to the electrochemical cell and had no impact current production (Data not shown). To confirm the effect of Antimycin A on biological current, addition of Antimycin A added to sterile minimal media in a reactor was performed and shown to have no impact on current production (**Fig. S4**)

#### Protein Collection and Quantification

Protein quantification was used to assess the total biomass in bio-electrochemical experiments. At the end of an electrochemical experiment the spent media from the reactor (∼20 mL) was collected and the biofilm was scraped from the working electrode. Biomass was centrifuged at 8,000 × *g* and the pellet was resuspended in 2mL of 10% w/v NaOH. Total protein collected from the reactor was quantified using a Qubit TM Protein Assay Kit (Invitrogen, USA) according to manufacturer’s protocol.

#### Statistical Methods

Uncertainties in bio-electrochemical measurements were calculated by analysis of at least three replicates for each bio-electrochemical experiment. All statistical analyses were performed in Excel and/or R.

Cathodic midpoint potentials were calculated from cyclic voltammogram (CV) scans. A full CV scan (between the aforementioned parameters found under electrochemical conditions) was separated into a forward scan (from 222 mV to −322 mV vs. SHE) and a reverse scan (−322 mV to 222 mV). The scan range was chosen to contain the voltage at which maximum current production was achieved. Because linearity could not be assumed in this data to generate a function for a first derivative analysis, an alternative method was used to analyze the data directly. A cubic smoothing spline (spar = 0.70) was applied to the current data in R to remove noise, but still capture the general trend. An approximate derivative was then taken from these values. From this approximate derivative, the maximum current produced, and its corresponding voltage potential were found for both the forward and reverse scans, which were then averaged to find the midpoint potential. The midpoint potential from each replicate was then pooled and averaged to obtain the reported value.

Anodic and cathodic currents were determined by chronoamperometry (CA). An average of the final 100 data points of each CA scan was taken to determine the average current achieved for each biological replicate. The replicates were then averaged to determine the average current produced for each strain. Biological current was determined by subtracting the average current post antimycin addition from average cathodic current prior to addition. To determine if the average current of any of the mutants was significantly different from the wild-type, the cumulative current data was compiled in Excel and then fitted to a linear model in order to perform a type II analysis of variance (ANOVA) test with R. Following the ANOVA test, anodic and cathodic currents for all mutants were compared with wild-type by Tukey”s honestly significant difference (HSD) test to determine if any significant difference existed between them.

#### Phylogenetic Analysis

Phylogenetic trees of relatives of the genes identified in this work were generated by search for homologs, homolog alignment, and tree assembly by a maximum likelihood method. Approximately 120 to 200 homologs for each gene identified in this work were identified with the top homolog hit program that interrogates the Integrated Microbial Genomes and Microbiomes Database (https://img.jgi.doe.gov/)^46,47^. Homolog sets for each gene were aligned with the Muscle aligner^48^ with default parameters.

Phylogenetic trees were generated for each homology set by a maximum likelihood method with RAxML 8.2.11^49^. 100 trees were generated for each set of homologs for bootstrapping. Tree images and taxonomic metadata (**Figs. S7** to **S11**) were generated and annotated using the interactive Tree of Life (iTOL) program^50^.

## End Notes

### Data Availability

The datasets generated during and analyzed during the current study are available from the corresponding authors (A.R. and B.B.) on reasonable request.

### Code Availability

The Macroscope Image Analyzer software is available at https://github.com/barstowlab/macroscope-image-analyzer.

### Materials & Correspondence

Correspondence and material requests should be addressed to A.R. or B.B.

## Author Contributions

Conceptualization, A.R. and B.B.; Methodology, A.R., M.B., and B.B.; Investigation, A.R. F.S., L.T., J.S., O.A., I.A., L.K., M.B., and B.B; Writing - Original Draft, A.R., L. T., F.S., and B.B.; Writing - Review & Editing, A.R., J.S., F.S., and B.B.; Funding Acquisition, A.R. and B.B.; Resources, A.R. and B.B.; Supervision, A.R. and B.B.; Data Curation, A.R., L. T., F.S., and B.B.; Visualization, A.R., F.S., L.T., J.S., and B.B; Formal Analysis, A.R., F.S., L.T., J.S., and B.B;

## Acknowledgements

We thank L. Jurgensen and S. Medin for programming assistance, B. Pian for experimental assistance, and A.M. Schmitz for review of this manuscript. This work was supported in the Barstow lab by a Career Award at the Scientific Interface from the Burroughs Welcome Fund, Princeton University startup funds, Cornell University startup funds, and by U.S. Department of Energy Biological and Environmental Research grant DE-SC0020179. In the Rowe lab, this work was funded by Air Force Office of Scientific Research grant FA9550-19-1-0305 and University of Cincinnati startup funds. The Baym lab is supported by an award from the David and Lucille Packard Foundation.

