## Supplementary Information for "Identification of a Pathway for Electron Uptake in *Shewanella oneidensis*"

### Supplementary Information Figures

**Fig. S1.** Time course of $AHDS_{red}$ oxidation by wild-type *S. oneidensis*.

**Fig. S2.** Representative time courses of anticipated hits from $AHDS_{red}$ oxidation screen of the *S. oneidensis* whole genome knockout collection.

**Fig. S3.** Representative time courses of unanticipated hits from $AHDS_{red}$ oxidation screen of the *S. oneidensis* whole genome knockout collection.

**Fig. S4.** Example electrochemical measurements of an *S. oneidensis* biofilm.

**Fig. S5.** $AHDS_{red}$ oxidation rates and biological cathodic currents produced by selected mutants of *S. oneidensis*.

**Fig. S6.** Aerobic to anaerobic and anaerobic to aerobic transitional growth curves of wild-type *S. oneidensis* mutants and selected mutants.

**Fig. S7.** Phylogenetic tree constructed for 120-200 of the closest identified genes to *SO_0841* in the Integrated Microbial Genes Database.

**Fig. S8.** Phylogenetic tree constructed for 120-200 of the closest identified genes to *SO_0181* in the Integrated Microbial Genes Database.

**Fig. S9.** Phylogenetic tree constructed for 120-200 of the closest identified genes to *SO_0400* in the Integrated Microbial Genes Database.

**Fig. S10.** Phylogenetic tree constructed for 120-200 of the closest identified genes to *SO_3660* in the Integrated Microbial Genes Database.

**Fig. S11.** Phylogenetic tree constructed for 120-200 of the closest identified genes to *SO_3662* in the Integrated Microbial Genes Database.

### Supplementary Information Tables

**Table S1.** Full results of $AHDS_{red}$ oxidation screens of the *S. oneidensis* whole genome knockout collection.

**Table S2.** Electrochemical data observed on cathodes for selected *S. oneidensis* transposon insertion mutants and controls.

**Table S3.** Growth and electrochemical data for deletion mutants of *SO_0181*, *SO_0400*, *SO_0841*, *SO_3660*, *SO_3662* and their corresponding complementation strains.

**Table S4.** Metadata for phylogenetic trees.

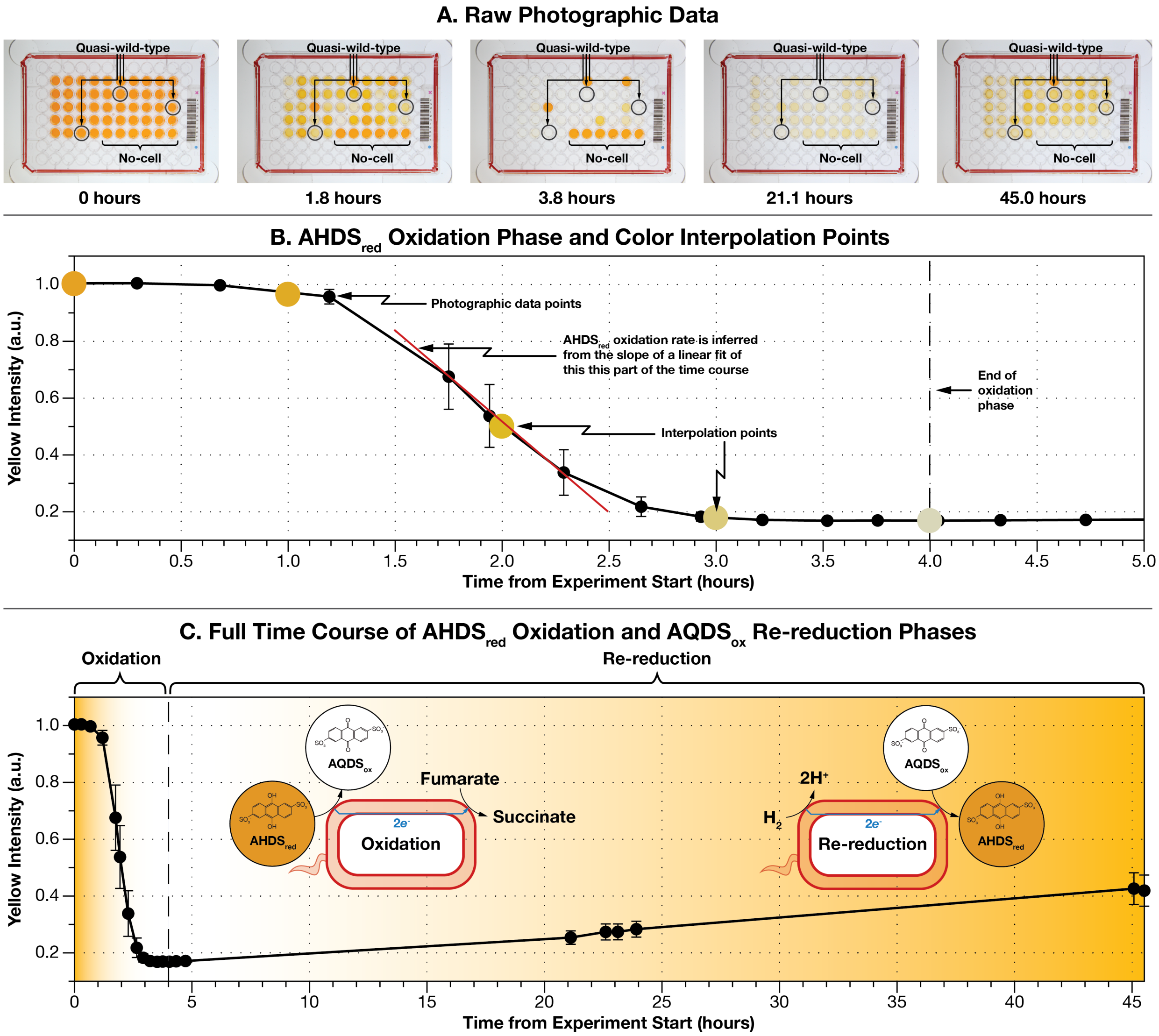


**Figure S1.** Time course of $AHDS_{red}$ oxidation by wild-type *S. oneidensis*. Quasi-wild-type mutants contain an transposon insertion but have no effect on $AHDS_{red}$ oxidation. (**A**) Raw photographic time-series of a single $AHDS_{red}$ oxidation assay plate. Mutants that behave like wild-type (quasi-wild-type) and no-cell controls are highlighted in the time-series of photographs. All wells in columns 1 and 12, and rows A, G and H are blank (no cells, and no $AHDS_{red}$/$AQDS_{ox}$). (**B**) Close up of the $AHDS_{red}$ oxidation phase (hours 0 to 4 from the start of the experiment) showing yellow intensity interpolation points used in the color graphs in **Fig. 1** in the main text and linear fit used to calculate $AHDS_{red}$ oxidation rates reported in **Fig. S5**. (**C**) Long time course of yellow intensity of average of quasi-wild-type wells showing $AHDS_{red}$ oxidation phase, and the subsequent re-reduction phase caused by transfer of electrons from $H_2$ in the headspace of the anaerobic chamber to $AQDS_{ox}$ mediated by *S. oneidensis*. Note that the no-cell controls slowly oxidize over ≈ 40 hours due to residual $O_2$ in the anaerobic chamber (< 20 ppm).

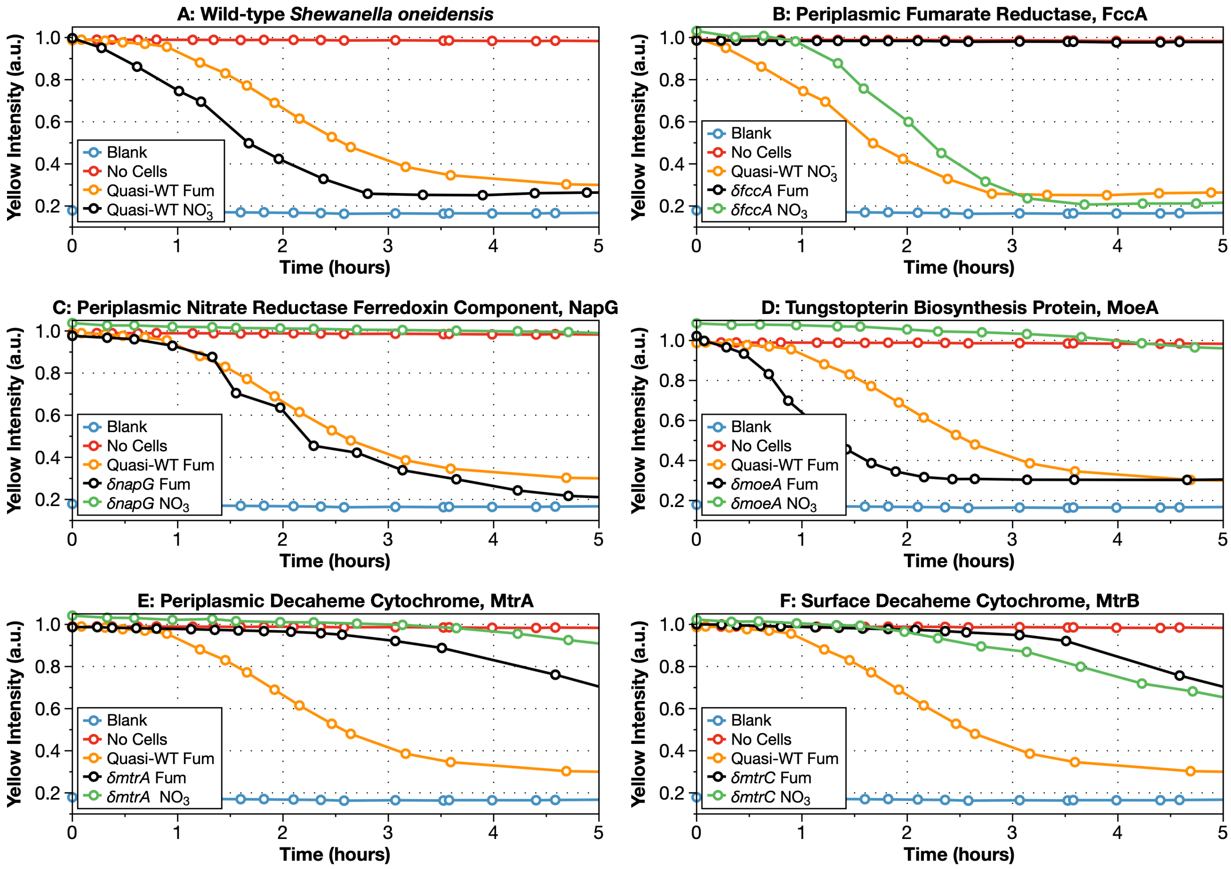


**Figure S2.** The $AHDS_{red}$ oxidation screen finds anticipated hits from the *S. oneidensis* whole genome knockout collection. (**A**) Quasi-wild-type *S. oneidensis* oxidizing $AHDS_{red}$ with fumarate (Fum) and nitrate ($NO_3^-$) terminal electron acceptors. (**B**) Disrupting the periplasmic fumarate reductase FccA knocks out $AHDS_{red}$ oxidation when fumarate is used as a terminal electron acceptor, but not nitrate. (**C** and **D**) Conversely, disruption of the nitrate reductase ferredoxin component (encoded by *napG*) or the MoeA enzyme that enables synthesis of it co-factor (coded by *moeA*) disrupts $AHDS_{red}$ oxidation when using nitrate as a terminal electron acceptor, but not fumarate. (**E** and **F**) Disrupting the MtrA and MtrC multi-heme cytochrome components of the Mtr EET complex slows AHDSred oxidation with both nitrate and fumarate.

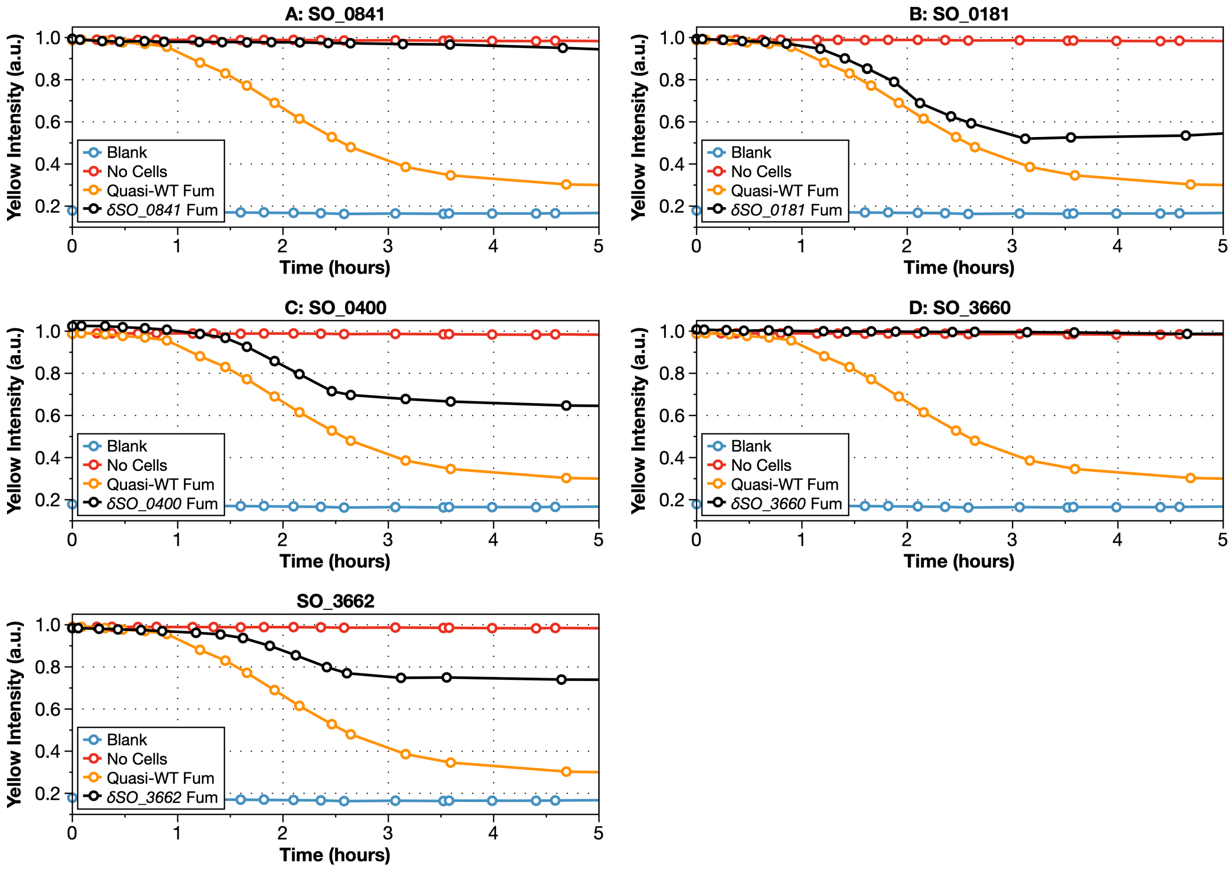


**Figure S3.** The $AHDS_{red}$ oxidation screen of the *S. oneidensis* whole genome knockout collection found 5 hits that produce robust disruption of electron uptake from a cathode. Here were show representative time courses of $AHDS_{red}$ oxidation coupled to fumarate reduction for disruption mutants of these 5 genes.

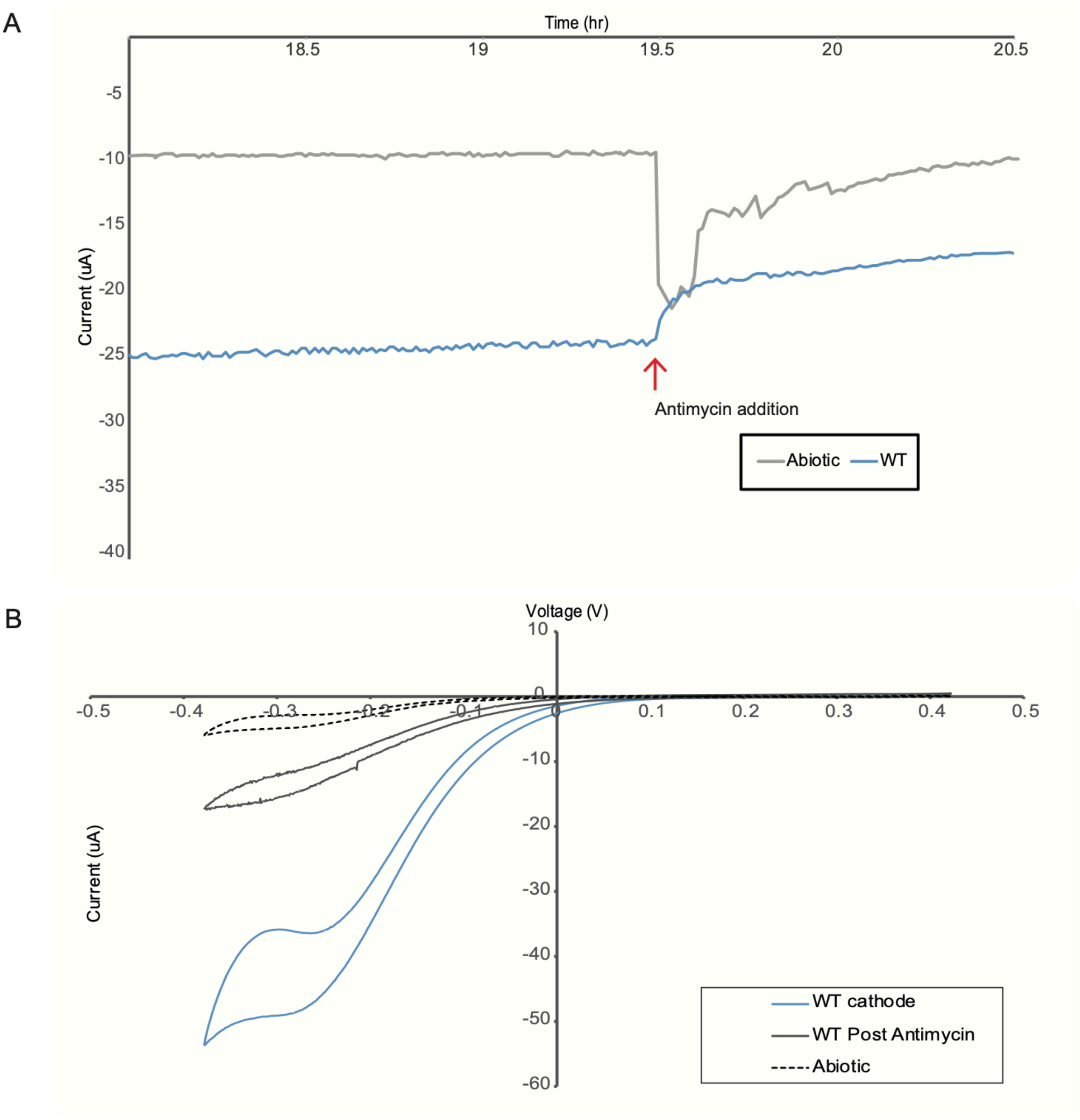


**Figure S4.** Example electrochemical measurements of an *S. oneidensis* biofilm. (**A**) Representative biological current production by wild-type *Shewanella oneidensis* MR-1 indicating total cathodic current and drop in current after the addition of the biological inhibitor Antimycin (50 µM) as compared with an abiotic control. (**B**) Cyclic voltammograms of the corresponding abiotic, cathodic and inhibited conditions for sample in panel **A**.

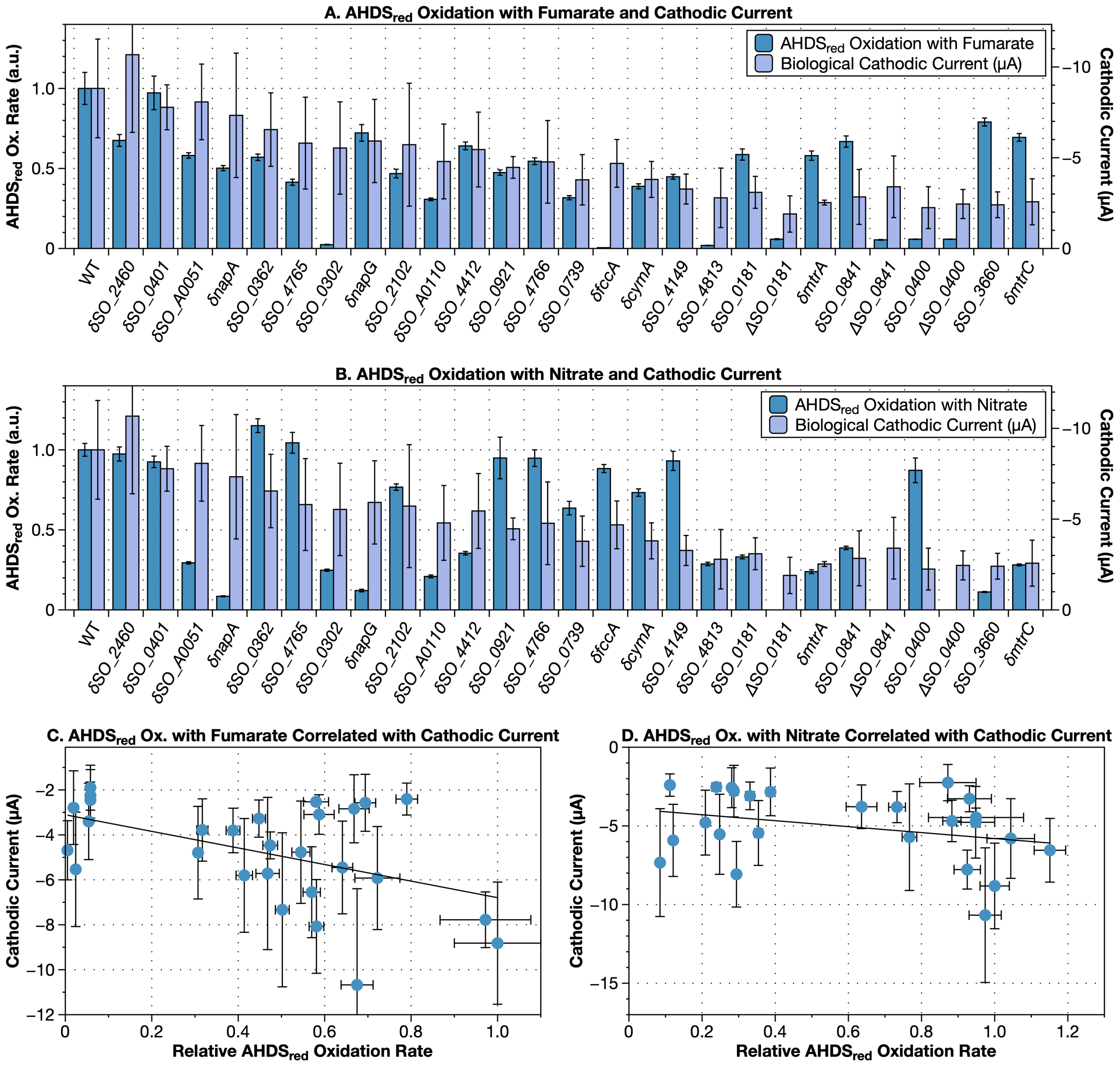


**Figure S5.** AHDS$_{red}$ oxidation rates and biological cathodic currents produced by selected mutants of *S. oneidensis.* Selected mutants are controls or produced AHDS$_{red}$ oxidation failure for unknown reasons. δ indicates a transposon insertion mutant, while Δ indicates a gene deletion mutant. The AHDS$_{red}$ oxidation rate is measured from the linear section of the yellow intensity trace as shown in **Fig. S1** and is reported relative to the wild-type oxidation rate. (**A**) AHDS$_{red}$ oxidation with fumarate and cathodic current. (**B**) AHDS$_{red}$ oxidation with nitrate and cathodic current. (**C**) AHDS$_{red}$ oxidation with fumarate correlated with cathodic current. (**D**) AHDS$_{red}$ oxidation with nitrate correlated with cathodic current.

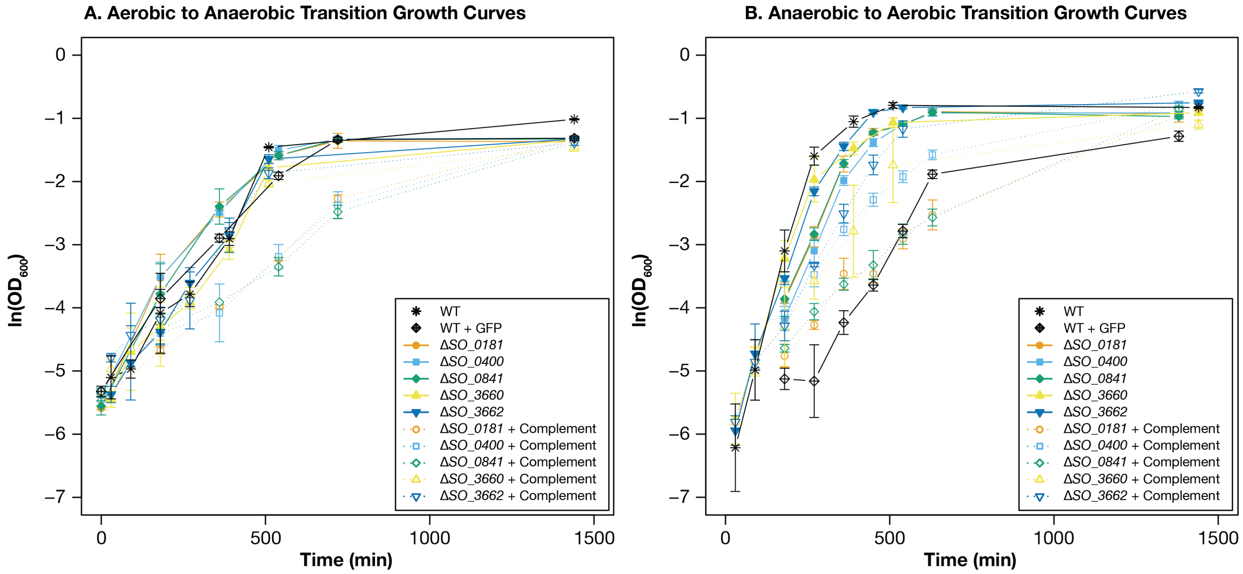


**Figure S6**. Aerobic to anaerobic and anaerobic to aerobic transitional growth curves of wild-type *S. oneidensis* mutants and selected mutants. (**A**) Aerobic to anaerobic transitional growth curves for clean deletion mutants and complements for genes *SO_0181*, *SO_0400*, *SO_0841*, *SO_3660*, and *SO_3662*, compared to wild-type. (**B**) Anaerobic to aerobic transitional growth curves for mutants and wild-type. Growth curves were measured in minimal media with 10 mM lactate as an electron donor and either oxygen or fumarate (20 mM) as an electron acceptor.

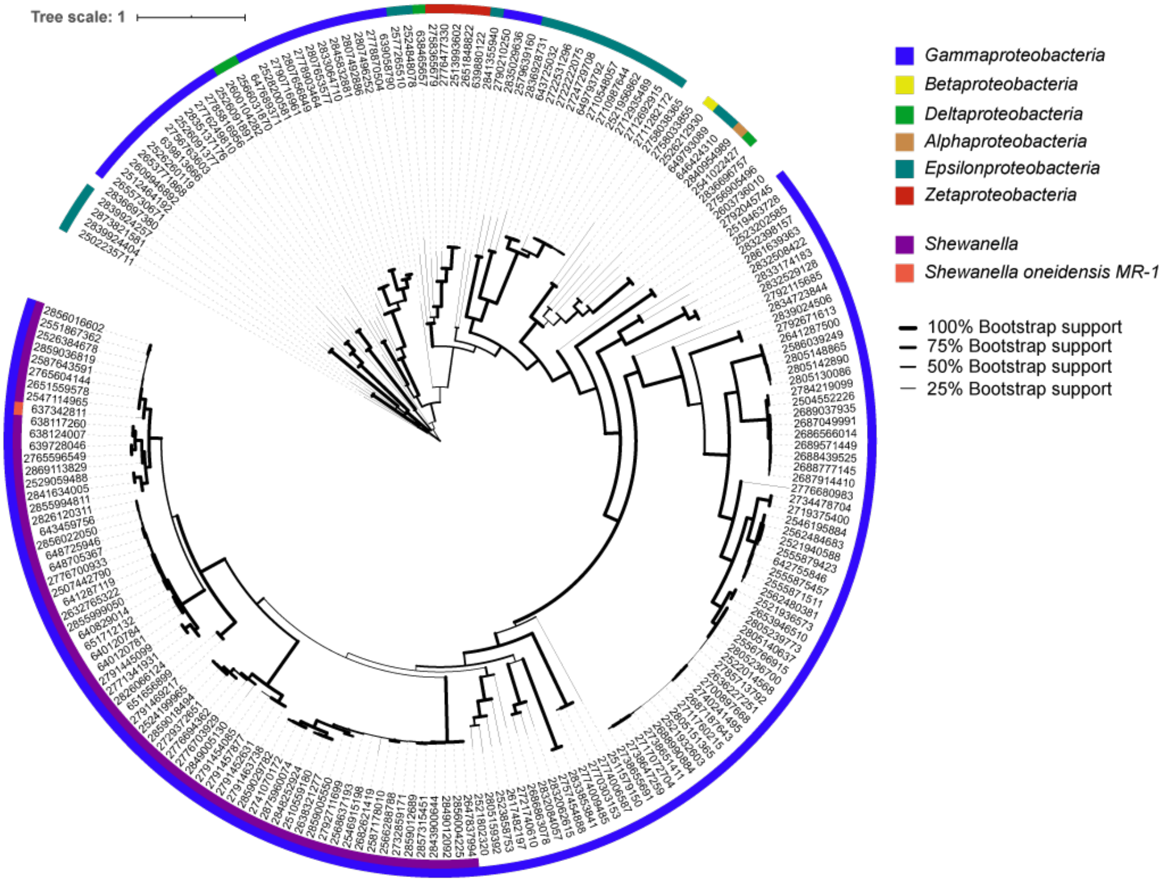


**Figure S7.** Phylogenetic tree constructed for 120-200 of the closest identified genes to *SO_0841* in the Integrated Microbial Genes Database (https://img.jgi.doe.gov/). Alignments generated using Muscle 3.8.425 using default parameters. A best scoring maximum likelihood tree was generated using RAxML 8.2.11 using 100 bootstrap replicates to identify the optimal tree. The tree was annotated using the interactive tree of life interface (https://itol.embl.de/). Thickness of branches indicates boot strap support for each branch. Color of outer-ring indicates phylum with a focus on *Proteobacteria*. Inner ring denotes homologs from *Shewanella* species with the strain identified in this study highlighted (*Shewanella oneidensis* MR-1). Metadata for trees attached in supplementary **Table S4**.

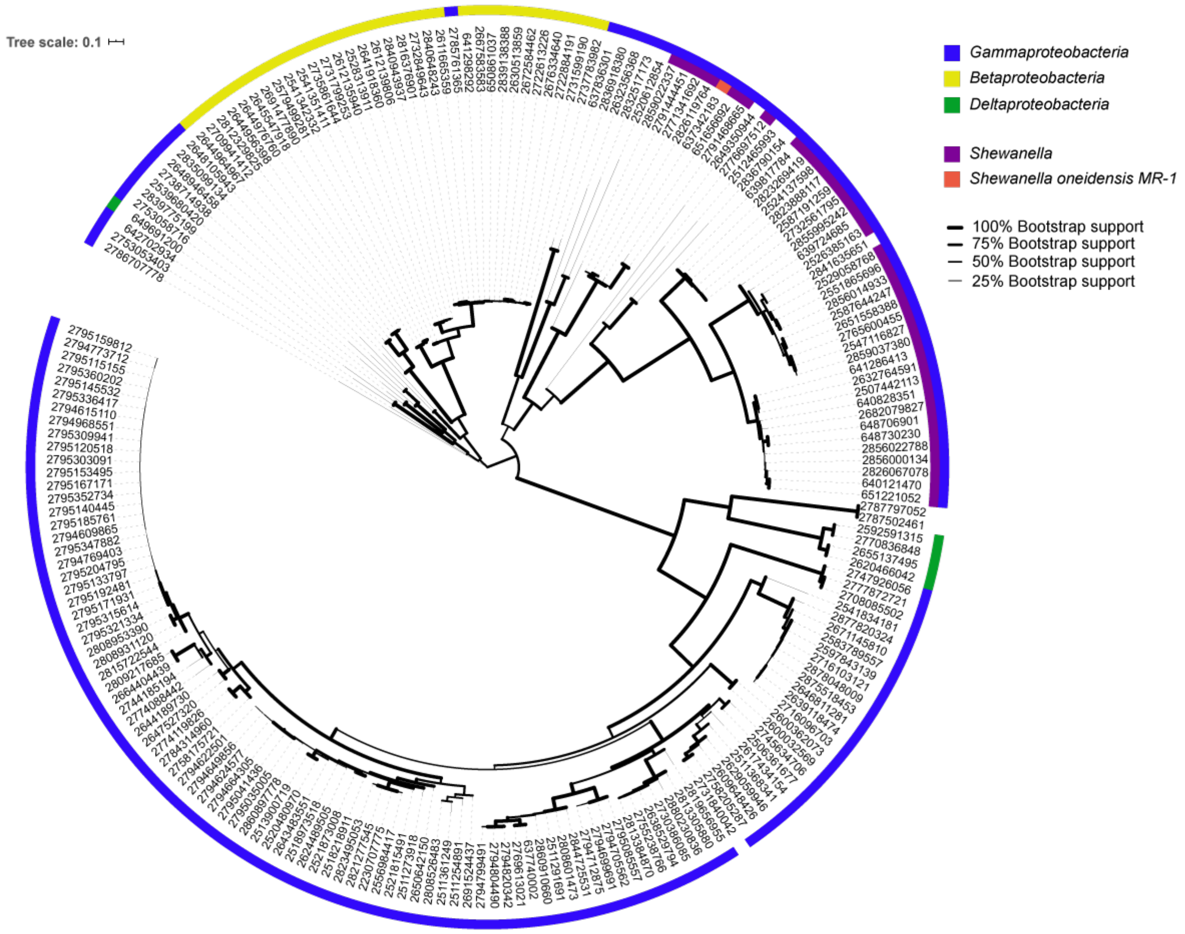


**Figure S8.** Phylogenetic tree constructed for 120-200 of the closest identified genes to *SO_0181* in the Integrated Microbial Genes Database (https://img.jgi.doe.gov/). Alignments generated using Muscle 3.8.425 using default parameters. A best scoring maximum likelihood tree was generated using RAxML 8.2.11 using 100 bootstrap replicates to identify the optimal tree. The tree was annotated using the interactive tree of life interface (https://itol.embl.de/). Thickness of branches indicates boot strap support for each branch. Color of outer-ring indicates phylum with a focus on *Proteobacteria*. Inner ring denotes homologs from *Shewanella* species with the strain identified in this study highlighted (*Shewanella oneidensis* MR-1). Metadata for trees attached in supplementary **Table S4**.

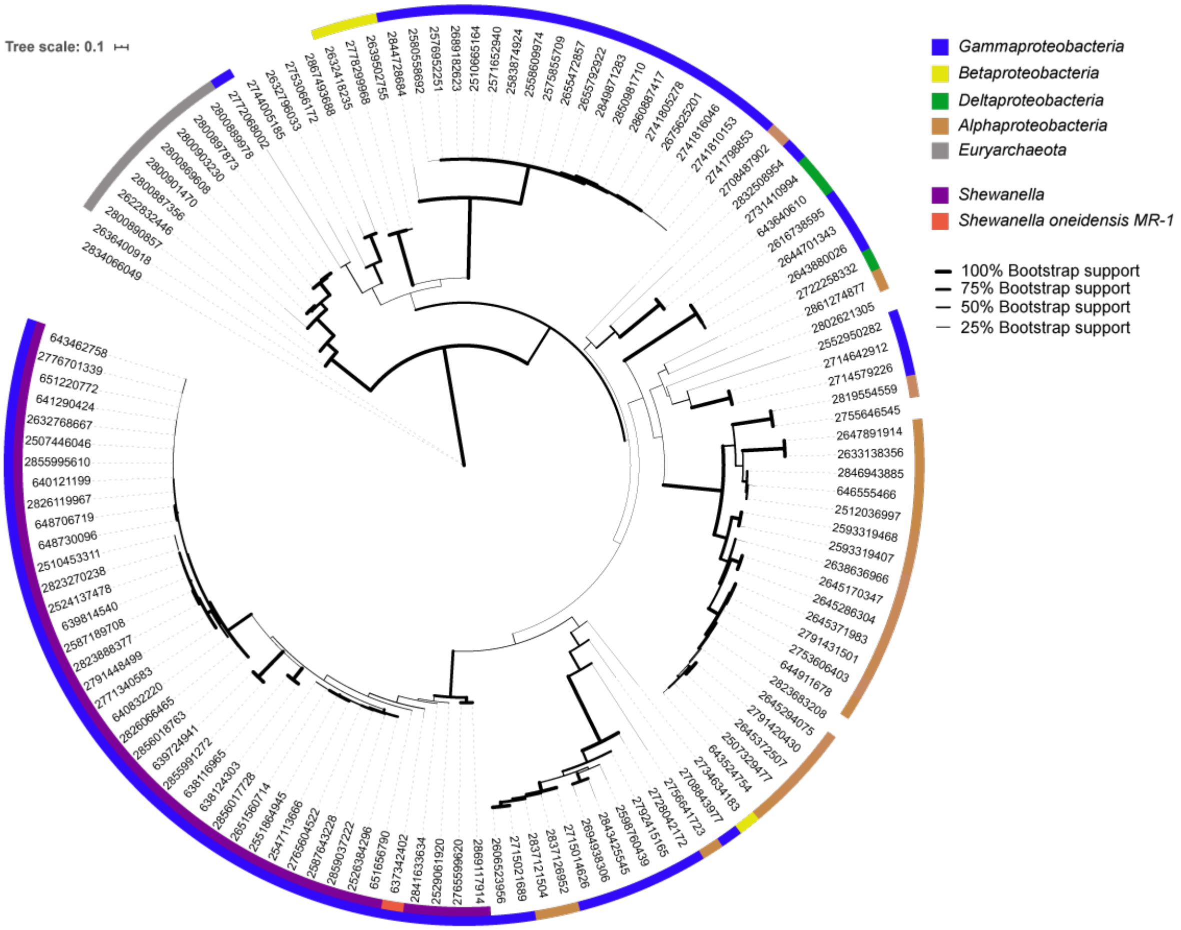


**Figure S9.** Phylogenetic tree constructed for 120-200 of the closest identified genes to *SO_0400* in the Integrated Microbial Genes Database (https://img.jgi.doe.gov/). Alignments generated using Muscle 3.8.425 using default parameters. A best scoring maximum likelihood tree was generated using RAxML 8.2.11 using 100 bootstrap replicates to identify the optimal tree. The tree was annotated using the interactive tree of life interface (https://itol.embl.de/). Thickness of branches indicates boot strap support for each branch. Color of outer-ring indicates phylum with a focus on *Proteobacteria*. Inner ring denotes homologs from *Shewanella* species with the strain identified in this study highlighted (*Shewanella oneidensis* MR-1). Metadata for trees attached in supplementary **Table S4**.

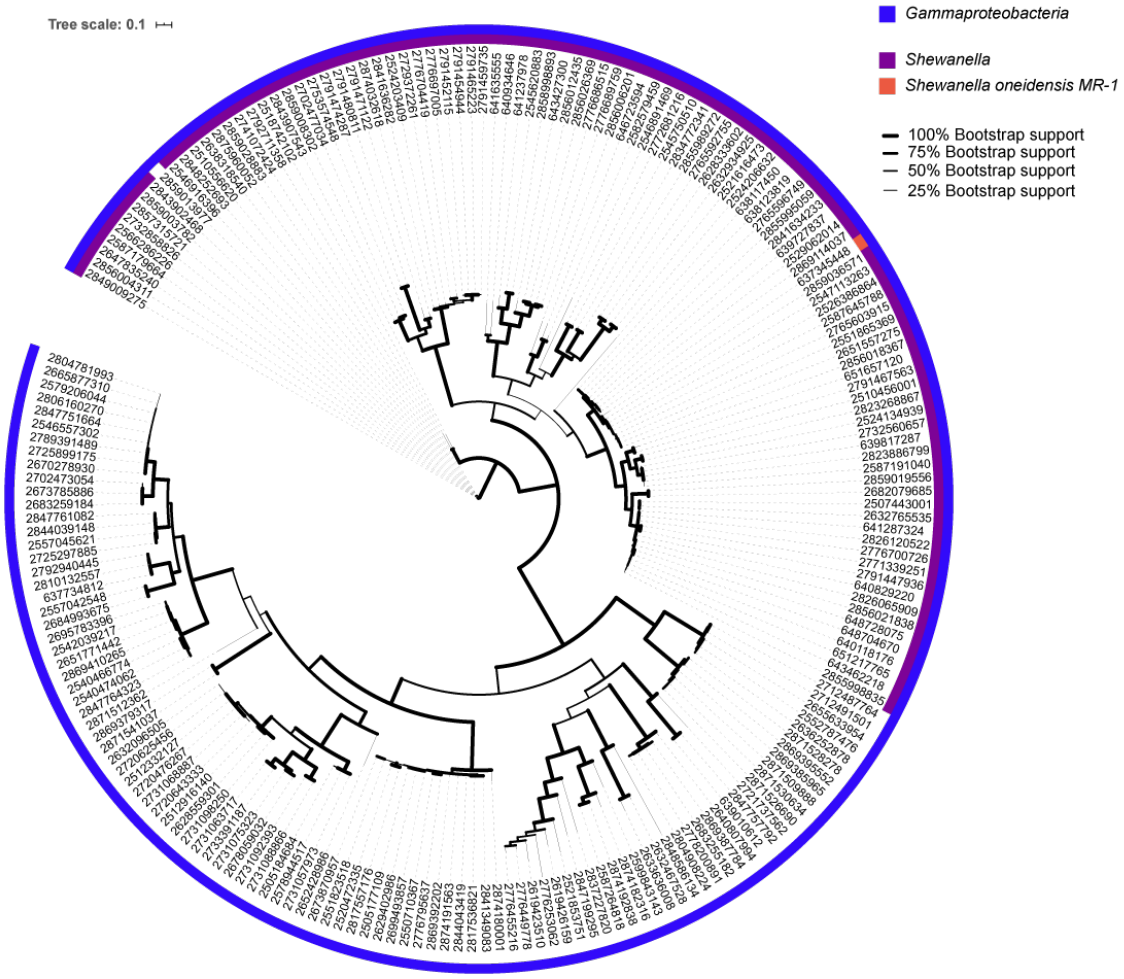


**Figure S10.** Phylogenetic tree constructed for 120-200 of the closest identified genes to *SO_3660* in the Integrated Microbial Genes Database (https://img.jgi.doe.gov/). Alignments generated using Muscle 3.8.425 using default parameters. A best scoring maximum likelihood tree was generated using RAxML 8.2.11 using 100 bootstrap replicates to identify the optimal tree. The tree was annotated using the interactive tree of life interface (https://itol.embl.de/). Thickness of branches indicates boot strap support for each branch. Color of outer-ring indicates phylum with a focus on *Proteobacteria*. Inner ring denotes homologs from *Shewanella* species with the strain identified in this study highlighted (*Shewanella oneidensis* MR-1). Metadata for trees attached in supplementary **Table S4**.

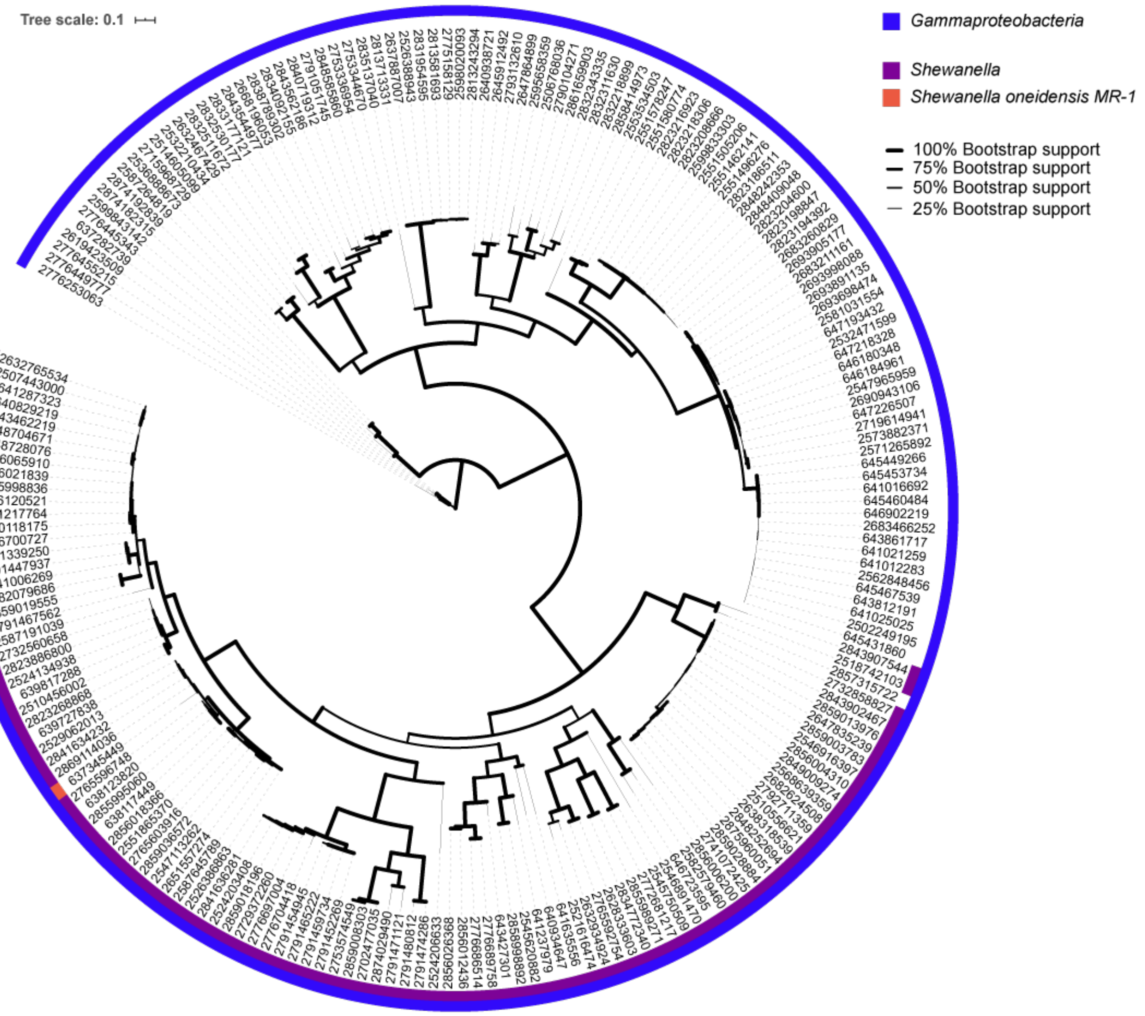


**Figure S11.** Phylogenetic tree constructed for 120-200 of the closest identified genes to *SO_3662* in the Integrated Microbial Genes Database (https://img.jgi.doe.gov/). Alignments generated using Muscle 3.8.425 using default parameters. A best scoring maximum likelihood tree was generated using RAxML 8.2.11 using 100 bootstrap replicates to identify the optimal tree. The tree was annotated using the interactive tree of life interface (https://itol.embl.de/). Thickness of branches indicates boot strap support for each branch. Color of outer-ring indicates phylum with a focus on *Proteobacteria*. Inner ring denotes homologs from *Shewanella* species with the strain identified in this study highlighted (*Shewanella oneidensis* MR-1). Metadata for trees attached in supplementary **Table S4**.

**Table S1.** A whole genome screen of the *S. oneidensis* whole genome knockout collection found 150 hits that disrupted $AHDS_{red}$ oxidation. The table shows the results of $AHDS_{red}$ oxidation screens with fumarate and nitrate terminal electron acceptors. A 'Y' in column H or I indicates that the mutant was either slow at oxidizing $AHDS_{red}$ or completely failed to do so with fumarate (column H) or nitrate (column I). A 'N' in either column indicates that the rate of $AHDS_{red}$ oxidation was not noticeably different from the wild-type oxidation rate for that terminal electron acceptor.

A 'Y*' in column H or I indicates that slowing or complete failure of $AHDS_{red}$ oxidation was observed but unreliable for that mutant and terminal electron acceptor. Typically, the mutant would only slow or eliminate $AHDS_{red}$ oxidation in ≈ 50% of experimental replicates. Several of these unreliable mutants were selected for electrochemical testing because of an interesting annotation. 'ND' in either column indicates that data for that combination of mutant and terminal electron acceptor is unavailable.

Disruption Location (column D) indicates the genomic coordinates of the transposon mutant for the locus. In some cases, multiple mutants that caused $AHDS_{red}$ oxidation slowing or failure were identified for a locus. Genomic coordinates between 1 (origin or replication of chromosome) and 4,969,811 refer to nucleotides in the *S. oneidensis* chromosome, and coordinates between 4,969,812 (origin of replication of megaplasmid) and 5,131,424 refer to nucleotides in the *S. oneidensis* megaplasmid. The nominal location column (column E) is used for spreadsheet sorting.

Fractional Distance (column F) indicates the distance of the transposon into the coding region of the locus. It is calculated by finding the distance between the first base of the start codon and the transposon insertion junction and dividing by the length of the gene.

| **Strain** | **Average Cathodic Current (μA)** | **Average Biological current (μA)** | **Average Anodic Current (μA)** | **Average Protein (μg/mL)** |
|---|---|---|---|---|
| Wild-type | -22.2 ± 4.0 | -8.8 ± 3.0 | 2.2 ± 0.2 | 143 ± 30 |
| *δSO_2460* | -26.4 ± 9.3 | -10.7 ± 5.2 | 1.7 ± 0.3 | 94 ± 18 |
| *δSO_0401* | -23.0 ± 3.9 | -7.8 ± 1.5 | 1.9 ± 0.1 | 70 ± 39 |
| *δSO_A0051* | -19.9 ± 3.2 | -8.1 ± 2.5 | 1.5 ± 0.3 | 90 ± 31 |
| *δnapA* | -25.5 ± 10.4 | -7.3 ± 4.2 | 1.5 ± 0.4 | 149 ± 12 |
| *δSO_0362* | -18.1 ± 3.8 | -6.6 ± 2.3 | 2.0 ± 0.7 | 113 ± 45 |
| *δSO_4765* | -20.5 ± 4.2 | -5.8 ± 3.1 | 1.9 ± 0.3 | 91 ± 36 |
| *δSO_0302* | -23.9 ± 3.7 | -5.5 ± 3.1 | 1.9 ± 0.2 | 115 ± 32 |
| *δnapG* | -18.9 ± 6.9 | -6.9 ± 3.1 | 1.7 ± 0.4 | 75 ± 39 |
| *δSO_2102* | -24.3 ± 6.2 | -5.7 ± 3.9 | 1.2 ± 0.4 | 118 ± 35 |
| *δSO_A0110* | -10.9 ± 1.7 | -4.8 ± 2.4 | 2.6 ± 1.0 | 109 ± 75 |
| *δSO_4412* | -20.4 ± 3.5 | -5.5 ± 2.3 | 1.1 ± 0.6 | 101 ± 36 |
| *δSO_0921* | -19.8 ± 1.9 | -4.5 ± 0.7 | 2.2 ± 0.6 | 96 ± 17 |
| *δSO_4766* | -13.1 ± 5.4 | -4.8 ± 2.6 | 1.4 ± 0.4 | 61 ± 18 |
| *δSO_0739* | -32.3 ± 8.1 | -3.8 ± 1.7 | 2.0 ± 0.6 | 102 ± 37 |
| *δfccA* | -21.8 ± 6.7 | -4.7 ± 4.7 | 1.7 ± 0.4 | 140 ± 62 |
| *δcymA* | -10.7 ± 0.9 | -3.8 ± 1.1 | 0.4 ± 0.1 | 70 ± 33 |
| *δSO_4149* | -21.8 ± 2.5 | -3.3 ± 1.0 | 1.9 ± 0.1 | 114 ± 27 |
| *δSO_4813* | -14.3 ± 3.8 | -2.8 ± 2.0 | 1.2 ± 0.3 | 81 ± 33 |
| *δSO_0181* | -11.7 ± 3.6 | -3.1 ± 1.0 | 1.7 ± 0.1 | 128 ± 55 |
| *δmtrA* | -15.6 ± 0.9 | -2.5 ± 0.2 | 1.1 ± 0.1 | 99 ± 18 |
| *δSO_0841* | -17.2 ± 5.1 | -2.8 ± 1.8 | 1.2 ± 0.1 | 119 ± 34 |
| *δSO_0400* | -9.2 ± 1.4 | -2.3 ± 1.4 | 1.9 ± 0.3 | 89 ± 55 |
| *δSO_3660* | -9.9 ± 0.8 | -2.6 ± 0.8 | 1.7 ± 0.3 | 81 ± 18 |
| *δmtrC* | -12.8 ± 2.4 | -2.9 ± 1.4 | 1.2 ± 0.2 | 120 ± 35 |

**Table S2.** Electrochemical data observed on cathodes for selected *S. oneidensis* transposon insertion mutants and controls. Averages are calculated from the mean of $n \geq 3$ replicates. Errors are calculated as ± one standard deviation of the mean.

| **Strain** | **Aerobic doubling time (± standard deviation)** | **Anaerobic doubling time (± standard deviation)** | **Average Anodic current (μA)** | **Average Cathodic Biological current (μA)** | **Average Midpoint potential (V)** |
|---|---|---|---|---|---|
| Wild-type | 0.98 ± 0.31 | 1.41 ± 0.26 | 2.2 ± 0.2 | -8.8 ± 3.0 | -0.214 ± 0.01 |
| Wild-type + GFP | 0.97 ± 0.24 | 1.25 ± 0.22 | *n.d.* | *n.d.* | *n.d.* |
| *ΔSO_0181* | 1.27 ± 0.25 | 1.15 ± 0.24 | 2.2 ± 0.1 | -1.9 ± 1.0 | -0.194 ± 0.01 |
| *ΔSO_0181* + Complement | 1.37 ± 0.21 | 1.95 ± 0.53 | 1.1 ± 0.1 | -11.6 ± 3.7 | -0.195 ± 0.03 |
| *ΔSO_0400* | 1.16 ± 0.44 | 1.39 ± 0.22 | 1.4 ± 0.8 | -2.45 ± 0.8 | -0.205± 0.01 |
| *ΔSO_0400* + Complement | 1.43 ± 0.17 | 1.93 ± 0.24 | 1.2 ± 0.1 | -8.2 ± 3.2 | -0.201 ± 0.02 |
| *ΔSO_0841* | 1.05 ± 0.44 | 1.71 ± 0.36 | 2.7 ± 0.2 | -3.4 ± 1.7 | -0.216 ± 0.01 |
| *ΔSO_0841* + Complement | 1.50 ± 0.22 | 2.02 ± 0.46 | 0.7 ± 0.1 | -8.3 ± 2.4 | -0.217 ± 0.01 |
| *ΔSO_3660* | 1.55 ± 0.28 | 1.93 ± 1.08 | 1.4 ± 0.7 | -2.9 ± 1.2 | -0.198 ± 0.03 |
| *ΔSO3660* + Complement | 1.57 ± 0.34 | 2.08 ± 0.42 | 0.7 ± 0.1 | -7.4 ± 2.2 | -0.217 ± 0.01 |
| *ΔSO3662* | 1.65 ± 0.52 | 1.88 ± 0.79 | 1.1 ± 0.1 | -2.64 ± 1.5 | -0.197 ± 0.01 |
| *ΔSO3662* + Complement | 1.60 ± 0.51 | 1.72 ± 0.35 | 0.7 ± 0.1 | -7.5 ± 1.3 | -0.206 ± 0.02 |

**Table S3.** Growth and electrochemical data for deletion mutants of *SO_0181*, *SO_0400*, *SO_0841*, *SO_3660*, *SO_3662* and their corresponding complementation strains. Averages are calculated from the mean of $n \geq 3$ replicates Errors are calculated as ± one standard deviation of the mean. *n.d.* = not detected.
